# Nonlinear processing of shape information in rat lateral extrastriate cortex

**DOI:** 10.1101/341792

**Authors:** Giulio Matteucci, Rosilari Bellacosa Marotti, Margherita Riggi, Federica B. Rosselli, Davide Zoccolan

## Abstract

In rodents, the progression of extrastriate areas located laterally to primary visual cortex (V1) has been assigned to a putative object-processing pathway (homologous to the primate ventral stream), based on anatomical considerations. Recently, we found functional support for such attribution (Tafazoli et al., 2017), by showing that this cortical progression is specialized for coding object identity despite view changes – the hallmark property of a ventral-like pathway. Here, we sought to clarify what computations are at the base of such specialization. To this aim, we performed multi-electrode recordings from V1 and laterolateral area LL (at the apex of the putative ventral-like hierarchy) of male adult rats, during the presentation of drifting gratings and noise movies. We found that the extent to which neuronal responses were entrained to the phase of the gratings sharply dropped from V1 to LL, along with the quality of the receptive fields inferred through reverse correlation. Concomitantly, the tendency of neurons to respond to different oriented gratings increased, while the sharpness of orientation tuning declined. Critically, these trends are consistent with the nonlinear summation of visual inputs that is expected to take place along the ventral stream, according to the predictions of hierarchical models of ventral computations and a meta-analysis of the monkey literature. This suggests an intriguing homology between the mechanisms responsible for building up shape selectivity and transformation tolerance in the visual cortex of primates and rodents, reasserting the potential of the latter as models to investigate ventral stream functions at the circuitry level.

**Significance Statement:** Despite the growing popularity of rodents as models of visual functions, it remains unclear whether their visual cortex contains specialized modules for processing shape information. To addresses this question, we compared how neuronal tuning evolves from rat primary visual cortex (V1) to a downstream visual cortical region (area LL) that previous work has implicated in shape processing. In our experiments, LL neurons displayed a stronger tendency to respond to drifting gratings with different orientations, while maintaining a sustained response across the whole duration of the drift cycle. These trends match the increased complexity of pattern selectivity and the augmented tolerance to stimulus translation found in monkey visual temporal cortex, thus revealing a homology between shape processing in rodents and primates.

## Introduction

In rodents, primary visual cortex (V1) is bordered by at least 9 extrastriate visual areas, whose involvement in dorsal and ventral stream processing (i.e., extraction of, respectively, motion and shape information) has become the object of intense investigation. To date, most studies on this front have been inspired by the routing of visual information in the specialized channels that, in primates, selectively relay low temporal/high spatial frequency signals to the ventral stream only, and high temporal/low spatial frequency content to both the ventral and dorsal streams (Maunsell et al., 1990; Ferrera et al., 1994). Inspired by a possible homology with primates, and by the existence of distinct V1 subpopulations making (receiving) target-specific projections with (from) downstream (upstream) areas (Gao et al., 2010; Matsui and Ohki, 2013; Glickfeld et al., 2013; Ji et al., 2015), several investigators have mapped mouse visual areas with drifting gratings of various spatiotemporal frequencies (Van den Bergh et al., 2010; Andermann et al., 2011; Marshel et al., 2011; Roth et al., 2012; Tohmi et al., 2014), finding the signature of dorsal processing in medial and parietal extrastriate cortex, but only scant evidence of ventral computations in lateral extrastriate cortex. This is not surprising, because the increase in the complexity of shape selectivity and tolerance to image variation that is the signature of ventral processing cannot be routed from upstream areas, but has to emerge as the result of local, non-linear integration of presynaptic inputs (DiCarlo et al., 2012).

To find the signature of such computations, we recently investigated how visual objects are represented along rat lateral extrastriate areas (Tafazoli et al., 2017). Our experiments showed that the neuronal populations along this progression become increasingly capable of supporting discrimination of visual objects despite transformations in their appearance (e.g., due to translation and scaling), with the largest transformation tolerance achieved in the most ventral region: the laterolateral area (LL). This trend closely matches the one found along the primate ventral pathway, thus supporting a central role of LL as the apex of the rat object-processing hierarchy. At the same time, studying cortical representations of visual objects has the disadvantage of relying on idiosyncratic choices of the stimulus set (the object conditions) and rather complex interpretive approaches (e.g., information theory and machine learning). This makes the experimental design hard to standardize, and often leads to conclusions that are only partially overlapping across studies. The assignment of rat lateral extrastriate cortex to the ventral stream is no exception, as shown by the much weaker specialization for object processing found in this region by earlier studies (Vermaercke et al., 2014; Vinken et al., 2016).

This prompted us to verify the higher rank of LL, relative to V1, in ventral processing, by designing an experiment that exploited the benefits of parametric stimuli, but, rather than focusing on spatiotemporal tuning, compared the two areas in terms of the nonlinearity of the stimulus-response relationship and the tendency of a neuron to be selective for different stimulus orientations. The first property was measured by the degree to which neuronal responses were entrained to the phase of drifting gratings, and by the extent to which the structure of the neuronal receptive fields could be inferred through reverse correlation. The second property was assessed by detecting the presence of multiple peaks in the orientation tuning curves and by measuring their sharpness.

Critically, all these properties followed trends of variations that are consistent with the nonlinear summation of visual inputs that is expected to take place along the ventral stream, according to the predictions derived from: 1) a meta-analysis of the monkey literature; 2) a conceptual model of hierarchical ventral computations (Riesenhuber and Poggio, 1999); and 3) a state-of-the-art deep convolutional network trained for image classification (Simonyan and Zisserman, 2014). As such, our results suggest a strong homology between the mechanisms that are responsible for supporting higher-order processing of shape information in rodents, primates and brain-inspired machine vision systems.

## Materials and methods

### Animal preparation and surgery

All animal procedures were in agreement with international and institutional standards for the care and use of animals in research and were approved by the Italian Ministry of Health: project N. DGSAF 22791-A, submitted on Sep. 7, 2015 and approved on Dec. 10, 2015 (approval N. 1254/ 2015-PR). 18 naïve Long-Evans male rats (Charles River Laboratories), with age 3-12 months and weight 300-700 grams, underwent extracellular recordings in either primary visual cortex (V1) or laterolateral (LL) extrastriate visual cortex. Each rat was anesthetized with an intraperitoneal (IP) injection of a solution of 0.3 mg/kg of fentanyl (Fentanest^®^, Pfizer) and 0.3 mg/kg of medetomidin (Domitor^®^, Orion Pharma). The level of anesthesia was monitored by checking the absence of tail, ear and hind paw reflex, as well as monitoring blood oxygenation, heart and respiratory rate through a pulse oximeter (Pulsesense-VET, Nonin). A constant flow of oxygen was delivered to the rat throughout the experiment to prevent hypoxia. A constant level of anesthesia was maintained through continuous IP infusion of the same anesthetic solution used for induction, but at a lower concentration (0.1 mg/kg/h Fentanyl and 0.1 g/kg/h Medetomidin), by means of a syringe pump (NE-500; New Era Pump Systems). Temperature was thermostatically kept at 37°C using a heating pad to prevent anesthesia-induced hypothermia.

After induction, the rat was secured to a stereotaxic apparatus (Narishige, SR-5R) in flat-skull orientation (i.e., with the surface of the skull parallel to the base of the stereotax) and, following a scalp incision, a craniotomy was performed over the target area in the left hemisphere (typically, a 2x2 mm^2^ window) and the dura was removed to allow the insertion of the electrode array. When targeting V1, the coordinates of penetration were ~6.5 mm posterior from bregma and ~4.5 mm left to the sagittal suture (i.e., AP 6.5, ML 4.5). When targeting LL, the coordinates were ~1 mm anterior from lambda and ~1 mm medial from the cranial ridge (i.e., AP 8, ML 5). Throughout the procedure, the rat eyes were protected from direct light and kept hydrated by repeated application of an ophthalmic ointment (Epigel^®^, Ceva Vetem).

Once the surgical procedure was completed, and prior to probe insertion, the stereotax was placed on a rotating platform and the rat’s left eye was covered with black, opaque tape, while the right eye (placed at 30 cm distance from the monitor) was immobilized using a metal eye-ring anchored to the stereotax. The platform was then rotated, so as to align the right eye with the center of the stimulus display and bring the binocular portion of its visual field to cover the left side of the display. During the recordings, the eye and cortex were periodically irrigated using saline solution to keep them properly hydrated.

### Neuronal recordings

Extracellular recordings were performed using single-shank 32-channel silicon probes (NeuroNexus Technologies) with site recording area of 775 μm^2^ and 25 μm of inter-site spacing. After grounding (by wiring the probe to the animal’s head skin), the electrode was manually lowered into the cortical tissue using an oil hydraulic micromanipulator (Narishige, MO-10; typical insertion speed: ~ 5 μm/s), up to the chosen insertion depth (~800-1,000 μm from the cortical surface when targeting V1, and ~2,500 μm when targeting LL). To reach LL, the probe was tilted of 30° relative to the vertical to the surface of the skull (i.e., relative to the vertical to the base of the stereotax; Fig. 1A), whereas, for V1 recordings, it was inserted either perpendicularly (about half of the sessions) or with a variable tilt, between 10° and 30° (remaining half of the sessions). Extracellular signals were acquired using a system three workstation (Tucker-Davis Technologies) with a sampling rate of 25 kHz. Before insertion, the probe was coated with Vybrant^®^ DiI cell-labeling solution (Invitrogen, Oregon, USA), to allow visualizing the probe insertion track *post-mortem*, through histological procedures. To this aim, at the end of the recording session, an electrolytic lesion was also performed by delivering current (5 μA for 2 seconds) through the 4 deepest channels at the tip of the shank.

### Visual stimuli

During a recording session, two kinds of visual stimulation protocols were administered to the rat.

A 15 min-long receptive field (RF) mapping procedure was used to precisely identify the visual area each neuron was recorded from (see details below) and optimize the location of the RFs for the main presentation protocol (i.e., ensure that most RFs fell inside the monitor, by rotating the platform or repositioning the eye through adjustments of the eye-ring). Following (Niell and Stryker, 2008), during the RF mapping protocol, the animal was presented with 10°-long drifting bars with four different orientations (0°, 45°, 90° and 135°), tiling a grid of 66 different visual field locations (i.e., 6 rows, spanning vertically 50°, and 11 columns, spanning horizontally 100°). The bars were white over a black background. During presentation of these stimuli, multiunit spiking responses (MUA) were plotted as a function of screen position in real-time, so as to estimate the positions of the RF centers across the recording sites of the probe. This allowed identifying in real-time when area LL was reached during an oblique probe insertion. Specifically, area LL was identified by the third “rightwards” (i.e., from nasal to temporal) reversal of the retinotopy at a recording depth close to 2.5 mm DV (Tafazoli et al., 2017). The same analysis was applied off-line to well-isolated single units obtained by spike sorting (see below) to precisely identify the neurons that were recorded from LL, as illustrated in Figure 1B.

Once the probe was positioned in the final recording location, the main presentation protocol was administered. The rat was presented with: 1) 1s-long drifting gratings, made of all possible combinations of 3 spatial frequencies (SF; 0.02, 0.04 and 0.08 cpd), 3 temporal frequencies (TF; 2, 4 and 8 Hz), and 12 directions (from 0° to 330°, in 30° increments); and 2) 30s-long movies of spatially and temporally correlated noise. To allow for a precise estimate of neuronal tuning properties, each grating stimulus was presented in 20 repeated trials, while 80 distinct noise movies were shown for an overall duration of 40 minutes. All stimulus conditions were randomly interleaved, with a 1s-long inter stimulus interval (ISI), during which the display was set to a uniform, middle-gray luminance level (later used to estimate spontaneous firing rates; see below). To generate the movies, random white noise movies were spatially correlated by convolving them with a Gaussian kernel. The kernel full width at half maximum (FWHM) was chosen to match a 0.02 cpd SF, which was found to elicit a robust spiking response in a series of pilot experiments. For the same reason (i.e., to increase the chance of evoking spikes), temporal correlations were also introduced, by convolving the movies with a causal exponential kernel with a 33 ms decay time-constant.

Stimuli were generated and controlled in MATLAB (The MathWorks) using the Psychophysics Toolbox package and displayed with gamma correction on a 47-inch LCD monitor (SHARP PNE471R) with 1920×1080 pixel resolution, 220 cd/m^2^ maximum brightness and spanning a visual angle of 110° azimuth and 60° elevation. Grating stimuli were presented at 60 Hz refresh rate, whereas noise movies were played at 30 Hz.

### Histology

At the end of the recording session, each animal was deeply anesthetized with an overdose of urethane (1.5 gr/kg) and perfused transcardially with phosphate buffer saline (PBS) 0.1 M, followed by 4% paraformaldehyde (PFA) in PBS 0.1 M, pH 7.4. The brain was then removed from the skull, post-fixed in 4% PFA for 24 h at 4°C, and then immersed in cryoprotectant solution (15% w/v sucrose in PBS 0.1 M, then 30% w/v sucrose in PBS 0.1 M) for 48 h at 4 °C. The brain was finally sectioned into 30μm-thick coronal slices using a freezing microtome (Leica SM2000R, Nussloch, Germany). Sections were mounted immediately on Superfrost Plus slides and let dry at room temperature overnight. A brief wash in distilled water was performed, to remove the excess of crystal salt sedimented on the slices, before inspecting them at the microscope. Each slice was then photographed with a digital camera adapted to a Leica microscope (Leica DM6000B-CTR6000, Nussloch, Germany), acquiring both a DiI fluorescence image (700 nm DiI filter) and a bright-field image at 2.5X and 10X magnification. Following the acquisition of this set of images, the sections displaying the electrode fluorescent track were further stained for Nissl substance using a Cresyl Violet Acetate solution, and new pictures were taken at 2.5X and 10X magnification. By superimposing the fluorescence, bright-field and Nissl-stained images, it was possible to reconstruct the tilt and the anteroposterior (AP) position of the probe during the recording session, as well as the laminar location of the recording sites. Specifically, the boundaries between the cortical layers were identified, based on the difference in size, morphology and density of the Nissl-labeled cells across the cortical thickness. The position of the probe relative to such boundaries was determined by tracing the outline of the fluorescent track, and taking into account, when available, the location of the electrolytic lesion performed at the end of the recording session. Based on the known geometry of the silicon probe, it was possible to draw the location of each recording site over the shank, thus estimating its laminar location. More details can be found in (Tafazoli et al., 2017).

### Selection of the single units included in the analyses

Responses of single units were isolated offline by applying the spike sorting package KlustaKwik-Phy (Rossant et al., 2016). Automated spike detection, feature extraction and expectation maximization (EM) clustering were followed by manual refinement of the sorting using a customized version of the “Kwik-GUI” interface. Specifically, the manual curation of the automatic output was performed by taking into consideration many features of the candidate clusters: 1) the distance between their centroids and their compactness in the space of the principal components of the waveforms (a key measure of goodness of spike isolation); 2) the shape of the auto- and cross-correlograms (important to decide whether to merge two clusters or not); 3) the variation, over time, of the principal component coefficients of the waveform (important to detect and take into account possible electrode drifts); and 4) the shape of the average waveform (to exclude, as artifacts, clearly non-physiological signals). Clusters suspected to contain a mixture of one or more single units were separated using the “reclustering” feature of the GUI (i.e., by rerunning the EM clustering algorithm on the spikes of these clusters only). At the end of the manual refinement step, only responsive, well-isolated single units, with reproducible firing across repeated stimulus presentations, were included in the dataset used to perform all subsequent analyses. Specifically, such units were defined by having: i) less than 0.5% of “rogue” spikes within 2 ms in their autocorrelogram (i.e., units displaying a clear refractory period); ii) a mean stimulus response firing rate of 2 spikes/s above baseline (i.e., responsive units); and iii) responsiveness in at least 30% of the trials in one or more stimulus conditions (i.e., stable isolation for at least 6 trials during the recording and reproducible stimulus-evoked response). We also applied an additional, very loose screening on the reproducibility of the firing rate across repeated stimulus presentations, by including only neurons with a mean correlation coefficient of the direction-tuning curve across trials not lower than 0.03. The average baseline (spontaneous) firing-rate of each well-isolated unit was computed by averaging its spiking activity over every ISI condition. These criteria led to the selection of 105 units in V1 and 104 units in LL.

### Analysis of the responses to drifting gratings

The response of a neuron to a drifting grating was computed by counting how many spikes the neuron fired (on average, across repeated presentations of the stimulus) during the whole duration of stimulus presentation, and then subtracting the spontaneous firing rate (see above). To quantify the tuning of a neuron for the orientation and direction of drifting gratings, we computed two standard metrics, the orientation and direction selectivity indexes (OSI and DSI), which are defined as: OSI = (*R*_pref_ − *R*_ortho_)/(*R*_pref_ + *R*_ortho_), and DSI = (*R*_pref_ −*R*_opposite_)/(*R*_pref_ + *R*_opposite_), where *R*_pref_ is the response of the neuron to the preferred direction, *R*_ortho_ is the response to the orthogonal direction, relative to the preferred one (i.e., *R*_ortho_ = *R*_pref_ + *π*⁄2), and *R*_opposite_ is the response to the opposite direction, relative to the preferred one (i.e., *R*_opposite_ = *R*_pref_ + *π*). Values close to one indicate very sharp tuning, whereas values close to zero are typical of untuned units.

The modulation of the spiking response to drifting gratings at the temporal frequency *f*_1_ of the stimulus was quantified by a modulation index (MI) adapted from (Wypych et al., 2012) and defined as:

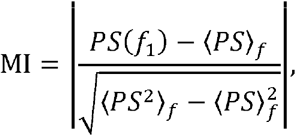
where *PS* indicates the power spectral density of the stimulus-evoked response, i.e., of the peri-stimulus time histogram (PSTH), and 〈 〉*_f_* denotes the average over frequencies. This metric measures the difference between the power of the response at the stimulus frequency and the average value of the power spectrum in units of its standard deviation. The power spectrum was computed by applying the Blackman-Tukey estimation method to the baseline-subtracted, 10ms-binned PSTH. Being MI a standardized measure, values greater than 3 can be interpreted as signaling a strong modulation of the firing rate at the stimulus frequency (typical of simple-like cells), whereas values smaller than 3 indicate poor modulation (typical of complex-like cells).

Bimodal tuning (i.e., the presence of two distinct peaks in the orientation-tuning curve) was quantified by the Bimodal Selectivity Index (David et al., 2006; Cadieu et al., 2007), defined as BSI = (*R*_2nd peak_ − *R*_2nd trough_) / (*R*_1st peak_ − *R*_1st trough_), where *R*_1st peak_ and *R*_2nd peak_ indicate the two largest peaks in the orientation-tuning curve (with *R*_1st peak_ > *R*_2nd peak_), and *R*_1st trough_ and *R*_2nd trough_ indicate the deepest troughs (with *R*_1st trough_^<^ *R*2nd trough). This index takes values close to one for tuning curves displaying two equally high peaks (regardless of their distance along the orientation tuning curve) and close to 0 for tuning curves with a single peak. Since this index is meant to provide a first-order assessment of the tuning for multiple oriented elements, it does not distinguish between cases where the orientation-tuning curve has two peaks from cases where it has more than two (i.e., the index discriminates curves with only one peak from curves with multiple peaks).

All the above metrics were computed using the responses of each neuron to the gratings shown at the most effective combination of spatial and temporal frequencies. For the vast majority of the recorded units in both areas (i.e., 97% and 86% in, respectively, V1 and LL), the preferred SFs were the lowest we tested (i.e., 0.02 and 0.04 cpd). Similarly, the preferred TFs were the lowest we tested (i.e., 2 and 4 Hz) for most neurons in both V1 (97%) and LL (88%).

### Analysis of the responses to the noise movies

We used the spike-triggered average (STA) analysis (Schwartz et al., 2006; Sharpee, 2013) to estimate the linear RF structure of each recorded neuron. The method was applied to the spike trains fired by the neuron in response to the spatiotemporally correlated noise (see above). The method yields an ordered sequence of images, each showing the linear approximation of the spatial RF at different times before spike generation (see examples in Fig. 2). To take into account the correlation structure of our stimulus ensemble and prevent artifactual blurring of the reconstructed filters, we “decorrelated” the resulting STA images by dividing them by the covariance matrix of the whole stimulus ensemble (Schwartz et al., 2006; Sharpee, 2013), using Tikhonov regularization to handle covariance matrix inversion. Statistical significance of the STA images was then assessed pixel wise, by applying the following permutation test. After randomly reshuffling the spike times, the STA analysis was repeated multiple times (*n* = 50) to derive a null distribution of intensity values for the case of no linear stimulus-spike relationship. This allowed z-scoring the actual STA intensity values using the mean and standard deviation of this null distribution (Figs. 2 and 4). The temporal span of the spatiotemporal linear kernel reconstructed via STA extended till 330 ms before spike generation, corresponding to a duration of 10 frames of the noise movie played at 30 Hz (Fig. 2). These procedures were performed on downsampled noise frames (16x32 pixels), with the resulting STA images that were later spline interpolated at higher resolution for better visualization and to obtain predictions of the neuronal responses using LN models (see next paragraph).

After estimating the linear spatiotemporal structure of a RF with STA, we used it as a filter in the input stage of a classical Linear-Nonlinear (LN) model of stimulus-response relationship (Schwartz et al., 2006; Sharpee, 2013). To obtain a prediction of the tuning of the neuron over the direction axis, the sequence of frames of each drifting grating was fed as an input to the STA-estimated linear filter. The output of the filter was then passed through a rectifying nonlinearity with unit gain to obtain the final response of the model to each stimulus frame. We finally integrated the response of the LN model through time to predict the activity of the neuron for each direction of the tuning curve. The agreement between the LN-predicted and observed tuning curves (black vs. colored curves in Fig. 2.i) was quantified by computing the fraction of variance explained by the model prediction. In addition, we also quantified how well the time course of the neuronal response to the preferred grating was explained by the time course of the LN-model prediction (black vs. colored curves in Fig. 2.ii, bottom), again, in terms of explained variance.

We also estimated the amount of signal contained in a given STA image by defining a quality metric based on a measure of maximal local contrast. Specifically, we used MATLAB “rangefilt” function (MATLAB Image Processing toolbox) to compute the local contrast in a STA image over convolutional patches with size equal to one fifth of the total image size and stride 1 (the local contrast in every patch was measured as the difference between the maximum and the minimum image intensity values within the patch itself). From the resulting local-contrast map (Fig. 5A, middle plot), we obtained a distribution of contrast intensity values (Fig. 5A, rightmost plot), and we defined a contrast index (CI) by taking the 0.9 quantile of this distribution (dashed line; the 0.9 quantile was chosen, instead of simply using the maximum local-contrast value, to make this measure more noise-robust). This CI metric was then used to quantify the amount of signal contained in the STA image. A STA-based RF was considered as well defined if, among the sequence of STA images corresponding to each pre-spike-generation time lag (see examples in Fig. 2), the one with the highest contrast had CI > 5.5 (usually, such highest-contrast image was the third in the sequence, corresponding to 66-99 ms before the spiking event; see examples in Fig. 5C). It should be noted that, since the intensity values of the original STA images were expressed as z-scores (see above), the 5.5 threshold can be interpreted in terms of peak-to-peak (i.e. white-to-black) distance in sigma units of the z-scored STA values. The 5.5 value was chosen in such a way to be conservative enough to include only STA images with a good signal-to-noise-ratio, but liberal enough to allow a fraction of LL neurons (whose STA images had typically poor contrast) to be considered for further analysis. This allowed a meaningful statistical comparison between V1 and LL in terms of lobe count (see next paragraph).

To further analyze the spatial structure of each well-defined, STA-based RF we automatically counted how many distinct lobes the highest-contrast STA image obtained for a unit contained. To this aim, the selected image was background-suppressed with the Perona-Malik algorithm (Perona and Malik, 1990) to enhance the high-intensity regions within the RF and reduce noise-driven false lobe detections. We then took the modulus of the resulting image and we set a threshold over its z-score values (ranging from 3 to 6 units of standard deviations; Fig. 5B). Based on such binarized image, we computed: i) the centroid position of each simply connected region (blue dots); and ii) the area-weighted center of mass (CM) of the image (green dot). Finally, we defined a “region of acceptability” for the centroids of the simply connected regions to be included in the lobe count. This region was a circle, centered on the CM, with a radius that was twice as large as the diameter of the largest simply connected region. Setting this acceptability circle allowed excluding spurious, simply connected (often very small) regions that were far away from the CM of the STA image (e.g., see the yellow dot in Fig. 5B), especially in the case of RFs with low CI. The total number of accepted centroids at the end of this selection procedure is the lobe count reported in Figure 5F for the different binarization thresholds.

### HMAX simulations

To obtain predictions about the evolution of the tuning properties of visual neurons along an object-processing pathway, we simulated ventral stream functional architecture using the HMAX model (Riesenhuber and Poggio, 1999). The structure of the model will be described at length in the Results (Fig. 3) and further motivated in the Discussion. Here, we provide some technical details about its specific implementation. For our application, we have chosen the version of the model described in (Serre et al., 2007) and downloaded from the website http://maxlab.neuro.georgetown.edu/docs/hmax/hmaxMatlab.tar. Briefly, the model is a feedforward neural network with alternating layers of units performing: 1) either a max-pooling (invariance-building) operation over input units with the same feature-selectivity, but RFs having different positions/scales (C1 and C2 units; dashed lines in Fig. 3); or 2) a template-matching (selectivity-building) operation over input units with different feature selectivity (S1 and S2 units; solid lines). As pointed out in the Discussion, the feature representations built by the model in the S-type layers are not learned, and there is no attempt to maximize classification accuracy in a given discrimination task. The S1 layer is simply a bank of Gabor filters with various orientations, spatial frequencies, positions and scales (similar to V1 simple cells), while the S2 units are tuned to random patches of images taken from various databases (e.g., Caltech 101), with different units having as a template the same image patch, but at different positions and scales. Such static, hardwired architecture makes the model very suitable to isolate the role of the max-pooling and template-matching computations in determining the tuning for oriented gratings along a ventral-like processing hierarchy.

To this aim, we fed as an input to the network (i.e., to the S1 layer) a set of drifting gratings spanning the same range of directions used in the neurophysiology experiment. Each drifting grating consisted of a sequence of progressively phase-shifted frames, and the activations of the units of the network in response to such frames were computed to simulate the dynamics of the neuronal responses during stimulus presentation. For each sampled unit, its preferred stimulus direction was selected and the power spectrum of its time-dependent (i.e., frame-dependent) response was computed to estimate the modulation index (MI), thus quantifying its phase sensitivity. We also integrated the activation of each unit over the whole stimulus presentation to compute its overall mean response for a given grating direction. By iterating this procedure for each stimulus direction, we built orientation and direction tuning curves and computed OSI and BSI values. All these indexes were computed for a randomly selected subset of units (*n* = 1000) in each layer, which was enough to obtain smooth distributions for the relevant indexes. The resulting distributions are those shown in Figures 4C, 7C and 8C.

### VGG16 simulations

We further checked whether the processing expected to take place along an object-processing hierarchy is consistent with the trends observed in rat visual cortex, by measuring the tuning for drifting gratins along the layers of VGG16, a state-of-the-art deep convolutional neuronal network (DCNN) for image classification (Simonyan and Zisserman, 2014). VGG16 is a large DCNN, totaling about 138 million parameters across its 16 convolutional and fully-connected layers. More specifically, the network consists in 5 blocks (colored boxes in Fig. 11A), made of 2 or 3 convolutional layers, with each block followed by a max-pooling layer (white boxes in Fig. 11A), and a final stack of three additional, fully-connected layers on top, before the softmax output layer for classification.

Detailed explanations of the roles the convolutional and pooling layers can be found elsewhere (LeCun et al., 2015). Briefly, the convolutional layer is an architectural prior that allows exploiting the structure of natural visual inputs (in particular, the spatial localization of visual objects and the ubiquity of translation as a commonly occurring identity-preserving transformation) to reduce the number of free parameters. Thanks to the structure of the visual world, a feature detector that is useful in one part of the visual field will likely be useful also in another part (because of the above-mentioned translation of visual features, caused by physical movements of objects and/or sensors). This allows “sharing” filter parameters across the visual field, by learning a single spatially localized filter in a given position and then applying it iteratively over the whole span of each input image. This also endows convolutional layers with spatial sparsity of connections: each unit in a convolutional feature map receives information only from a small, localized, subset of units in the previous layer (largely cutting down the number of parameters, as compared to a fully connected network). The pooling layer is another architectural prior that consists in performing a downsampling operation over the input (usually via a “max” computation, equivalent to the one implemented by HMAX), thus hardwiring in the output some amount of translation tolerance. This, again, can be seen as a way of leveraging on the prior knowledge that naturally occurring translation of visual objects preserve their identity, in order to shrink the width of the feature maps (thus reducing the number of parameters needed in the subsequent layers) and, at the same time, helping the network to build robust, invariant representations.

For our tests, we used VGG16 pre-trained as it was for the ILSVRC-2014 competition (Simonyan and Zisserman, 2014). The weights were downloaded from http://www.robots.ox.ac.uk/~vgg/research/very_deep/ via Keras (Chollet, 2015, GitHub, https://github.com/fchollet/keras) interface. Those weights were originally obtained by training the network on the ImageNet dataset – a large photographic database very popular as a computer vision benchmark (including over 14 million hand-annotated pictures with labels indicating which objects are pictured in each image). As done with HMAX, we fed to the input layer of the network a set of drifting gratings, resulting from combining 30 spatial frequencies, 4 temporal frequencies and 24 different directions. We then computed the activation (over the entire duration of each presented grating) of a pool of 1000 randomly selected units from the first convolutional layer of each block, from which we measured the relevant tuning properties of the units, to obtain the statistical characterization shown in Figure 11B and C.

### Experimental design and statistical analysis

To decide how many neurons to record in each of the two areas (V1 and LL) under investigation, we took inspiration from previous studies of rodent and monkey visual cortex (see Tables 1 and 2). In most of these studies, a good statistical assessment of the tuning properties of visual neurons (e.g., orientation tuning and phase sensitivity) was based on a sample size of > 100 units, although a few studies used less (~50 or fewer) and some used more (> 200). We thus set as a target for our study to collect at least 100 very well-isolate single-units (SUs) per area. Since the number of SUs obtained from each recording session was highly unpredictable, we performed multiple recording sessions from the same or different animals until our target number was reached in both areas. This required performing neuronal recordings from V1 of 12 different rats (for a total of 15 recording sessions) and from area LL of 6 different rats (for a total of 9 recording sessions), yielding a total of 105 and 104 responsive and reproducibly driven SUs in, respectively, V1 and LL. In V1, 5 sessions yielded more than 10 SUs, 4 sessions yielded between 5 and 10 SUs, while the remaining 6 sessions yielded less than 5 SUs. In LL, 5 sessions yielded more than 10 SUs, 3 sessions yielded between 5 and 10 SUs, while the last session yielded less than 5 SUs. When considered in terms of neurons yielded by each animal, the mean number of units recorded per rat was 8.75 in V1 and 17.33 in LL, while the minimal and maximal numbers of units per rat were, respectively, 1 and 29 in V1, and 3 and 45 in LL.

Throughout the study, differences between median values of two distributions were quantified using a Mann-Whitney U-test. Differences in the fraction of units above or below a certain threshold index value (for MI, OSI, BSI or CI) were quantified using the χ^2^ test for homogeneity with 1 degree of freedom. When needed (i.e., in case of small sample size), Fisher exact test was used to compare two distributions (e.g., the lobe count distributions in Fig. 5F). To check whether the distributions of the relevant indexes (i.e., MI, OSI, and BSI) were different between V1 and LL we applied a Kolmogorov-Smirnov test. To compare the average RF size extracted from the STA images, we applied an unpaired, one-tailed t-test with 65 degrees of freedom. To compare the information conveyed by neuronal responses about stimulus orientation in V1 and LL, we applied an unpaired, two-tailed, t-test with 207 degrees of freedom. To assess the significance of the correlation between the information about stimulus orientation and OSI, we used paired, two-tailed, t-tests with 103 (in V1) and 102 (in LL) degrees of freedom after applying Fisher transformation. Similarly, when testing for correlation between BSI and OSI, we adopted paired, two-tailed, t-tests with 103 (in V1) and 102 (in LL) degrees of freedom after applying Fisher transformation. Finally, in the meta-analysis shown in Figures 4D and 7D, we statistically compared the mean fractions of modulated (Fig. 4D) and orientation-tuned (Fig. 7D) units across primate visual cortical areas V1, V4 and IT using a one-way ANOVA with, respectively, 13 and 14 degrees of freedom.

## Results

We used 32-channel silicon probes to perform extracellular recordings from areas V1 and LL (Fig. 1A) of anaesthetized rats that were presented with drifting gratings of various orientations, spatial (SF) and temporal (TF) frequencies, as well as with 80 30s-long movies of spatially and temporally correlated noise (Materials and Methods). A total of 168 and 208 well-isolated single units were recorded, respectively, from V1 and LL. Among these neurons, 63% (i.e., 105 V1 units) and 50% (i.e., 104 LL units) were effectively and reproducibly driven by one or more of the grating stimuli across repeated presentations (Materials and Methods). Before the main stimulus presentation protocol, we also run a receptive field mapping procedure (Materials and Methods) to track the progression of the RFs recorded along a probe (Fig. 1B). Given that, in rodents, the polarity of the retinotopic map reverses at the boundary between each pair of adjacent visual cortical areas (Glickfeld and Olsen, 2017), tracking the retinotopy of the RFs allowed a precise identification of the units that were recorded from LL. This was possible because, to reach LL, the probe was inserted obliquely (with an angle of 30°, relative to the vertical to the surface of the skull), in such a way to first cross the lateral extrastriate areas that are medial to LL – i.e., the lateromedial (LM) and laterointermediate (LI) areas (Fig. 1A). In the case of V1 recordings, the probe was inserted either perpendicularly or with a variable angle (between 10° and 30°).

For each recording session, we also performed the histological reconstruction of the probe insertion track, appositely coated with a fluorescent dye before the penetration (see Fig. 1A). The outline of the probe, with the known location of the recording sites, was then superimposed to an image of the cortical section stained for Nissl substance, in such a way to infer the laminar location of the recorded single units (Material and Methods). In both areas, our recordings targeted the infragranular layers, because the tilted insertion that was necessary to track the retinotopy of lateral areas, thus allowing a reliable identification of LL (see previous paragraph), naturally brought the recording sites to land mostly in layer V. Therefore, in order to allow a fair statistical comparison with LL, the probe was aimed at the infragranular laminae also in V1. Our histological analysis confirmed that the laminar sampling was highly consistent between the two areas, with the vast majority of neurons being recoded from layer V in both V1 and LL (Fig. 1C). Critically, this ensures that the tuning properties being compared in our study refer to nearly identical laminar populations. At the same time, the selective targeting of layer V does not undermine the generality of our comparison. In fact, our previous investigation of V1 and lateral extrastriate areas showed that the same increase in the specialization for ventral processing along the areas’ progression was observable across the whole cortical thickness, and was equally sharp in superficial and deep layers (Tafazoli et al., 2017).

**Figure 1.**
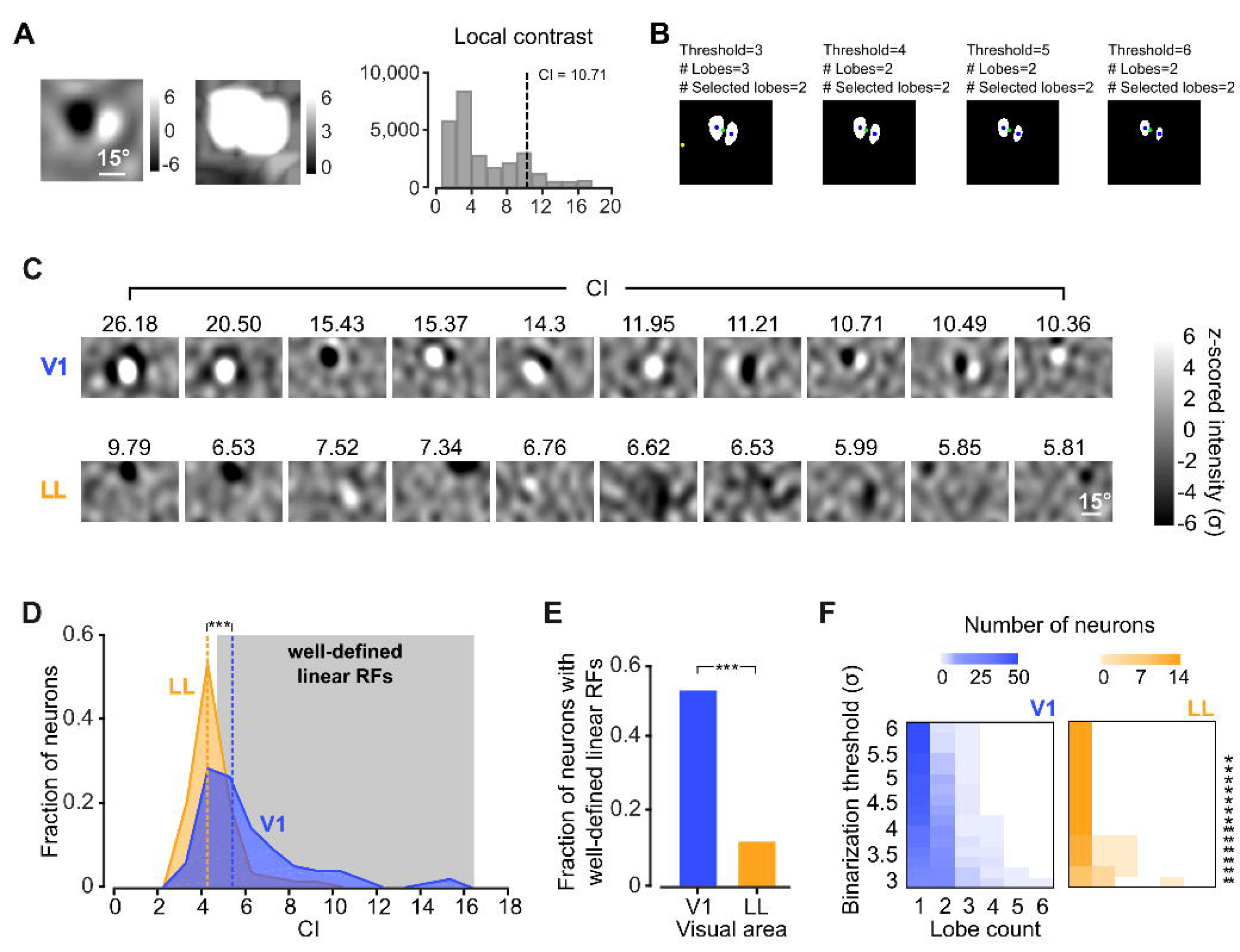
Targeting rat laterolateral (LL) visual cortical area for multi-electrode recordings using a linear silicon probe. **A**. Representative Nissl-stained coronal slice showing the insertion track of a single-shank silicon probe aimed at area LL. The probe was inserted obliquely into visual cortex, crossing lateromedial (LM) and laterointermediate (LI) visual areas before reaching LL. The dashed line depicts the border between LI and LL, as reconstructed by tracking the retinotopy of the receptive fields (RFs) recorded along the probe (see panel **B**). The position of the probe during the recordings is marked by the staining with the fluorescent dye (red) that was applied to it before insertion, and by the mechanical lesion produced by the penetration. Based on this information, it was possible to estimate the geometry of the probe (white lines) at the final recording depth (2.5 mm DV), along with the location of the recording sites (white dots), from the tip (site #1) to the base (site #32). **B**. Firing intensity maps displaying the RFs of the units recorded along the probe shown in **A**. The numbers refer to the sites each unit was recorded from. The reversal of the retinotopic progression (from leftward to rightward) at sites #13-14 marks the boundary between areas LI and LL. **C**. Laminar distributions of the responsive and reproducibly-driven V1 (blue) and LL (orange) single units, for which it was possible to establish the cortical layer through histological analysis.

### Examples of tuning curves and spatiotemporal receptive fields in V1 and LL

In both V1 and LL, neuronal responses to drifting gratings displayed a variety of tuning profiles that matched well the RF structures inferred through STA. This is illustrated in Figure 2, which reports, for each of two representative V1 (A-B) and LL (C-D) neurons: i) the tuning curve as a function of the direction of the grating, when presented at the most effective SF and TF; ii) the modulation of the firing rate elicited by the grating at the preferred direction; and iii) the sequence of STA images showing the spatiotemporal evolution of the RF structure. Each panel also reports the values of the metrics used to quantify the tuning properties of the neurons (Materials and Methods). Orientation and direction tuning were measured using the *orientation* and *direction selectivity indexes* (OSI and DSI), whose values range from 1 (maximal selectivity) to 0 (no selectivity), depending on the difference between the responses to the preferred orientation/direction and the orthogonal/opposite one. We also computed a *bimodal selectivity index* (BSI) to assess whether the tuning curve for orientation had multiple peaks. This index ranges from 0 (perfectly unimodal curve) to 1 (equal responsiveness to two non-adjacent orientations), and can be taken as a measure of complexity of tuning, with large BSI indicating selectivity for multiple oriented features (David et al., 2006; Cadieu et al., 2007). Finally, we computed a *modulation index* (MI) to quantify the extent to which the firing rate of a neuron was entrained to the phase of the gratings (Movshon et al., 1978; Skottun et al., 1991; Wypych et al., 2012), with MI > 3 indicating responses that were strongly phase-locked to the stimulus temporal frequency. As for the spatiotemporal filter estimated via STA, we used it as the linear stage of a Linear-Nonlinear (LN) model of stimulus-response mapping (Schwartz et al., 2006; Sharpee, 2013) to predict the tuning curve of the neuron over the direction axis and the time course of its response to the preferred grating (in both cases, prediction accuracy was quantified as the fraction of response variance that was explained by the model).

Figure 2A shows an example V1 neuron with sharp orientation tuning, good direction selectivity, and a response that was strongly modulated at the temporal frequency (4 Hz) of the preferred grating (blue curves/dots). These properties match those of a *simple cell* (Hubel and Wiesel, 1968), detecting the presence of an oriented edge at a specific visual field location – hence, the modulation of the firing rate, produced by the phasic alternation of light and dark oriented stripes, while the preferred grating swept across the neuron’s RF. Such position-dependent neuronal firing suggests that the response of the neuron to a visual pattern can be approximated as the sum of its responses to the individual constituent elements (the “pixels”) of the stimulus. Accordingly, the STA-reconstructed RF (Fig. 2A.iii) had a Gabor-like, double-lobed appearance, with an orientation matching the one corresponding to the peak of the tuning curve (Fig. 2A.i). When used as a filter in a LN model, the STA-based RF accurately predicted the tuning of the neuron over the direction and time axes (Figs. 2A.i and ii, respectively; black curves), thus confirming the linearity of the stimulus-response relationship. By contrast, the example unit shown in Figure 2B, while still being selective for a single orientation, displayed a highly non-linear behavior, which is typical of the class of neurons known as *complex cells* (Hubel and Wiesel, 1968). This is shown by the lack of modulation of the response to the preferred grating (Fig. 2B.ii), by the poorly defined spatiotemporal structure of the STA-based RF (Fig. 2B.iii), and by the failure of the latter to account for the tuning and time-modulation of the response (Fig. 2B.i-ii; compare blue and black curves).

**Figure 2.**
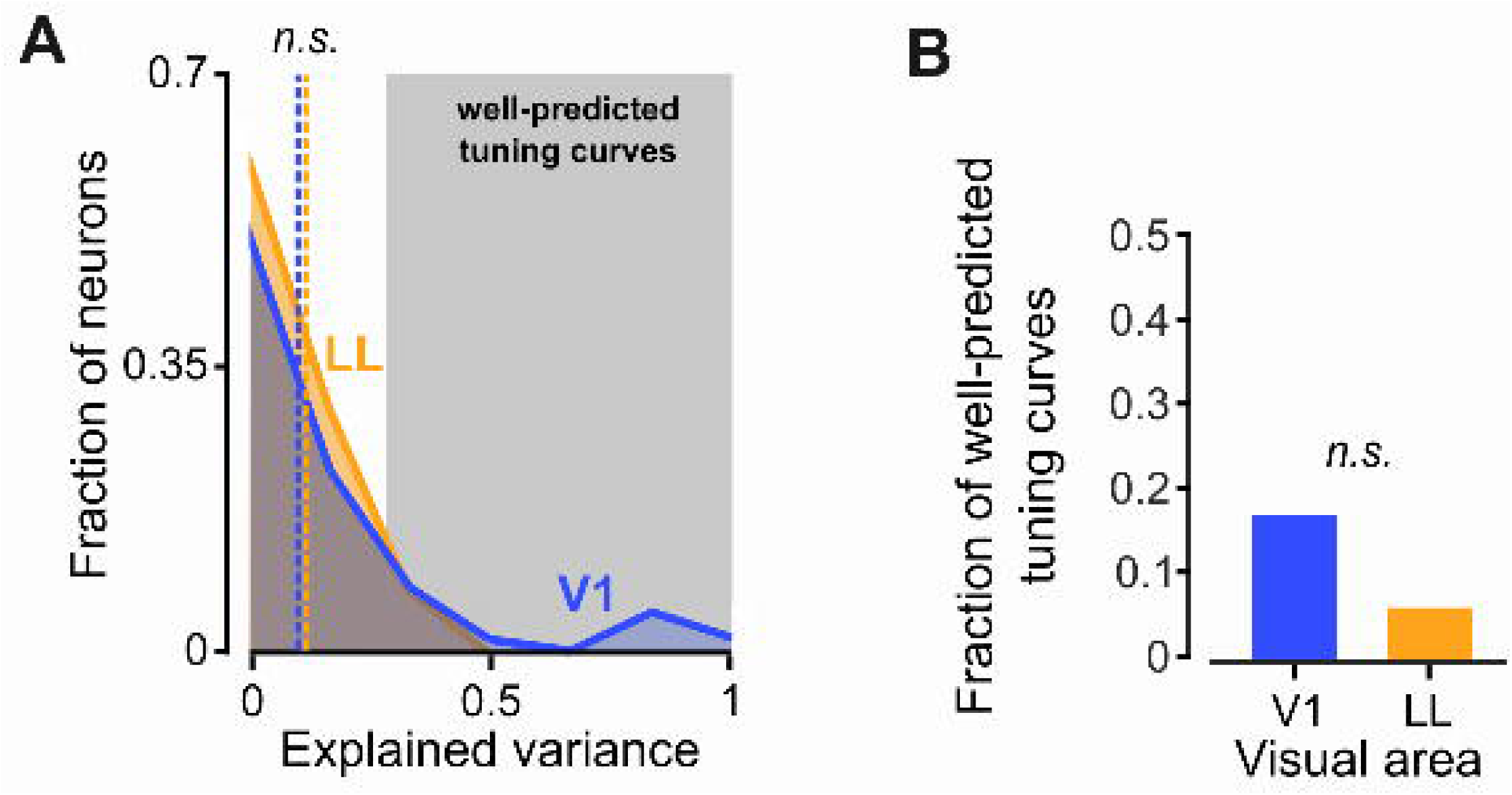
Examples of neuronal tuning in V1 and LL. Representative V1 simple (**A**) and complex (**B**) cells are shown in the upper panels (blue lines). Representative LL simple-like (**C**) and complex-like (**D**) cells are shown in the lower panels (orange lines). In each panel, subpanels i-iii illustrate different aspects of neuronal tuning. (i) The polar plot reports the observed (colored curve) and predicted (black curve) tuning of the neuron for the direction of a drifting grating, when shown at the preferred spatial (SF) and temporal (TF) frequencies of the unit. Predictions were obtained using a linear-nonlinear (LN) model, derived from the spike-triggered average (STA) images shown in iii. For ease of comparison, both the observed and predicted curves were normalized to their maximum. The plot also reports the fraction of variance of the tuning curve that was explained by the LN model, as well as the values of several metrics that quantify neuronal tuning: the orientation (OSI), direction (DSI) and bimodal (BSI) selectivity indexes. (ii) Raster plot (top) and peri-stimulus time histogram (PSTH; bottom; colored curve), showing the time course of the neuronal response to a drifting grating, when presented at the preferred SF, TF and direction. The bottom plot also reports the time course of the response, as predicted by the LN model (black curve), along with the fraction of variance explained by the model. For ease of comparison, both the observed and LN-predicted responses were normalized to their maximum. The top plot reports the value of the modulation index (MI), which measured the extent to which the neuronal response was time-locked to the phase of the drifting grating. (iii) Spatial structure and time-evolution of the STA-estimated RF of the neuron. In every STA image, each pixel intensity value was independently z-scored, based on the null distribution of STA values obtained through a permutation test (Materials and Methods). As such, all RF intensity values are reported as distances (in units of standard deviation σ; gray scale bar in panel D.iii) from what expected in the case of no frame-related, linear information carried by the spikes.

Interestingly, simple-like and complex-like units were also found in area LL. For instance, Figure 2C shows an LL neuron with a simple-like behavior, as indicated by the phase modulation of its response (Fig. 2C.ii; orange dots and curve), which was predicted with reasonable accuracy (black line) by the STA-reconstructed RF. This is consistent with the well-defined spatiotemporal structure of the RF (Fig. 2C.iii), which, however, was not able to explain as much variance of the direction-tuning curve (Fig. 2C.i; black vs. orange curve) as in the case of the example V1 simple cell (Fig. 2A.i). This could be due to the sensitivity of the neuron to multiple stimulus orientations, as shown by the large BSI value (0.52), which reflects the presence of two peaks (at 0° and 240°) in the direction-tuning curve (Fig. 2C.i; orange curve) – a behavior that cannot be captured by a RF with a linear, Gabor-like structure. Figure 2D shows instead an example LL unit that was sharply tuned for a single orientation, but displayed a typical complex-like behavior: its response to the preferred grating was unmodulated (Fig. 2D.ii), and both the direction-tuning curve (Fig. 2D.i) and the response dynamics (Fig. 2D.ii) were poorly explained by the STA-reconstructed RF (orange vs. black curves), which, in turn, lacked any clear spatiotemporal structure (Fig. 2D.iii).

These examples illustrate how it is possible to compare the level of linearity of the stimulus-response relationship for the neuronal populations recorded in V1 and LL, along with the sharpness and complexity (e.g., bimodality) of their tuning for orientation. To guide such comparison, we derived predictions about the evolution of neuronal tuning along an object processing hierarchy, by assessing how our grating stimuli were represented in HMAX – a neural network model of the ventral stream that is able to account for several qualitative trends in the tuning properties of ventral neurons (Riesenhuber and Poggio, 1999; Serre et al., 2007; Cadieu et al., 2007; Zoccolan et al., 2007; Cox and Riesenhuber, 2015; Leibo et al., 2017). In its simplest implementation, HMAX is a 4-layered, feed-forward network, with each layer consisting of either simple-like (S) or complex-like (C) units (Fig. 3). The first layer (S1) is a bank of Gabor filters that simulate V1 simple cells. In the second layer (C1), each unit performs a max pooling over a set of S1 afferents with the same orientation tuning, but slightly different RF positions or sizes. This yields orientation-tuned C1 units that are similar to V1 complex cells – i.e., they have larger RFs (dashed circles in Fig. 3), with increased position- and size-tolerance, as compared to their S1 afferents. In the third layer (S2), each unit performs a non-linear template-matching computation over a set of C1 afferent units with different preferred orientations. As a result, the S2 units are tuned for complex visual patterns, made of multiple oriented features, like the neurons in higher-level stages of the primate ventral stream, such as cortical area V4 and the inferotemporal cortex (IT) (DiCarlo et al., 2012). Finally, in the fourth layer (C2), each unit performs again a max pooling over S2 afferents that are tuned for the same visual pattern, but with slightly different RF positions and sizes. This results in C2 units with complex shape selectivity and large tolerance to position and size changes. Here, we used this basic 4-layered architecture to predict how the build-up of shape tuning and invariance should affect neuronal sensitivity to the orientation and temporal frequency of the gratings. In our simulations, layers S1 and C1 correspond to the simple and complex cells of V1, while layers S2 and, above all, C2, correspond to a downstream, higher-level area (such as LL).

**Figure 3.**
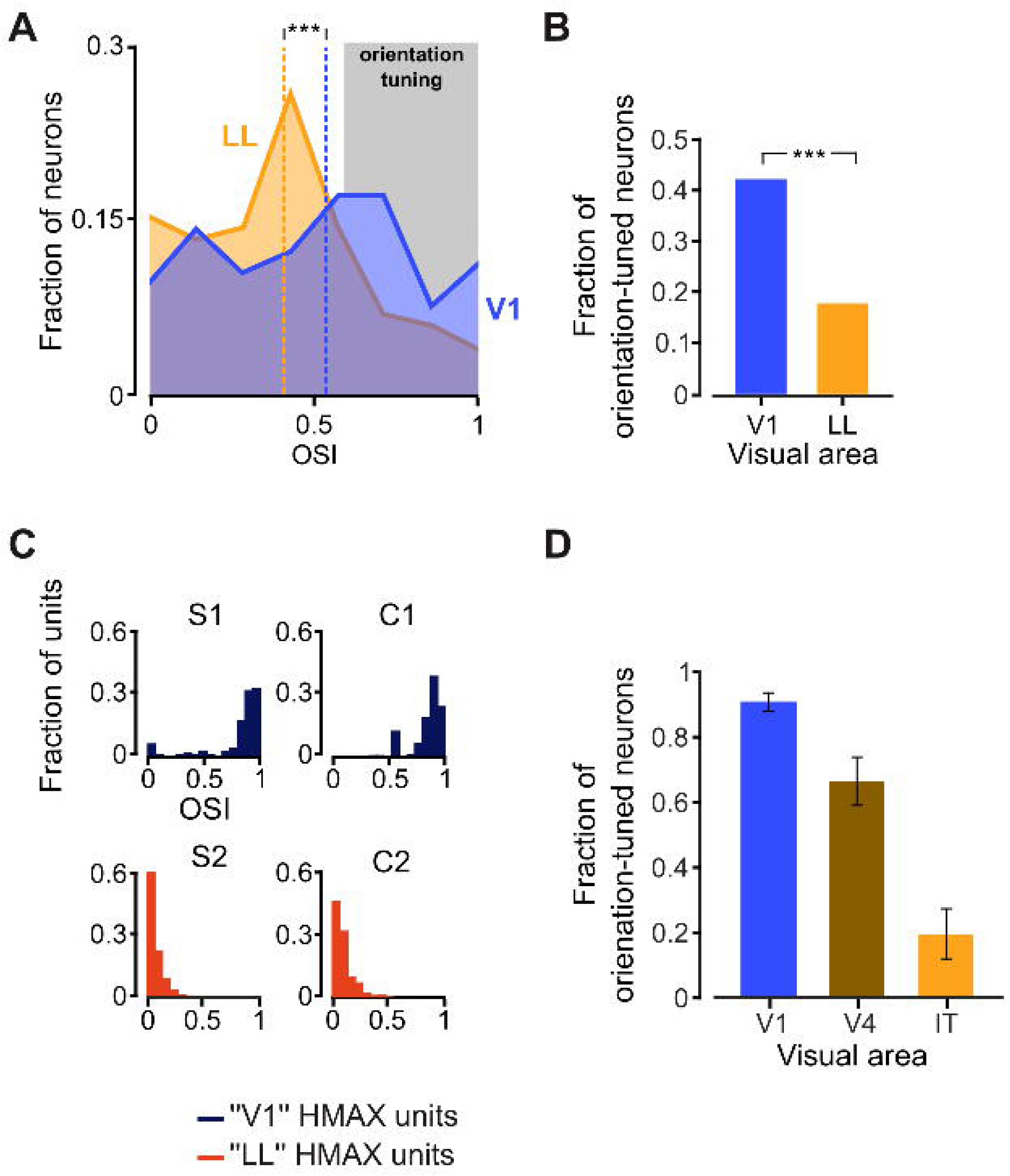
Sketch of the HMAX model implementation used in this study. The model is a feedforward, 4-layered neural network that processes arbitrary input images through a cascade of computations. The first layer (S1) consists of a bank of Gabor filters performing local edge detection on small input image patches (dark gray circles), similar to V1 simple cells. In the second layer (C2), each unit pools over many iso-oriented S1 units via a max-like operation (dashed lines), thus gaining some amount of position and scale tolerance, similar to V1 complex cells (such expansion of the invariance field is represented by the light gray rings). In the third layer (S2), each unit acquires tuning for more complex visual patterns by combining the output of many C1 units with different orientation via a non-linear template-matching operation (solid lines). In the fourth layer, each C2 unit applies again the max-pooling operation to S2 units with the same shape tuning but different RF positions and sizes, in order to further gain tolerance to such transformations. In our application, the input images were drifting gratings, similar to those used in the neurophysiology experiment, so as to obtain predictions about the evolution of the relevant tuning indexes (MI, OSI and BSI) across consecutive stages of a ventral-like hierarchy.

### The linearity of the stimulus-response relationship decreases from V1 to LL

To establish whether V1 and LL neurons differed in terms of the linearity of the stimulus-response relationship, we first compared the extent to which neuronal responses in the two areas were modulated by the phase of the drifting gratings. Despite the presence of both strongly phase-modulated and unmodulated units in both areas, the distributions of the modulation index (MI) were strikingly different in V1 and LL (*p* = 5.85*10^−6^; Kolmogorov-Smirnov test; Fig. 4A). In agreement with earlier studies of rat (Girman et al., 1999) and mouse (Niell and Stryker, 2008; Van den Bergh et al., 2010) primary visual cortex, the MI distribution obtained in V1 (blue curve) revealed the existence of two roughly equi-populated classes of neurons, falling, respectively, below and above the MI = 3 threshold that distinguishes complex-like from simple-like cells. By contrast, in LL, most MI values were lower than 3, following a distribution (orange curve) with a prominent peak at MI < 1, which is indicative of a strong prevalence of complex-like units. As a result, the median MI was much lower in LL than in V1 (1.28 vs. 2.55; orange vs. blue line) and this difference was highly significant (*p* = 8.04*10^−6^; Mann-Whitney U-test). Similarly, the fraction of simple-like cells in LL was half as large as in V1 (Fig. 4B; *p* = 1.93*10^−4^, χ^2^ test, *df* = 1).

As illustrated in Figure 2D.ii, the poor sensitivity to the phase of the drifting gratings observed in LL indicates that most neurons in this area respond to phasic stimuli with a sustained firing. This suggests that, in LL, most neurons are tolerant to position changes of their preferred visual stimuli. This interpretation was supported by the evolution of the modulation index observed in HMAX (Fig. 4C), where phase-sensitive cells were only found in the bottom layer (S1), corresponding to V1 simple cells. By contrast, from the C1 layer onward, all units displayed a fully phase-invariant behavior, because of the nonlinear max pooling that, in C1, implements position and size invariance. As a result, the combined distribution of MI values obtained in S1 and C1 was in qualitative agreement with the mixture of simple and complex cells observed in V1 (compare to Fig. 4A; blue curve). Similarly, the extreme phase invariance observed in S2 and C2 was qualitatively consistent with the stark prevalence of complex-like cells found in LL (> 80%; compare to Fig. 4A and B; orange curve/bar).

**Figure 4.**
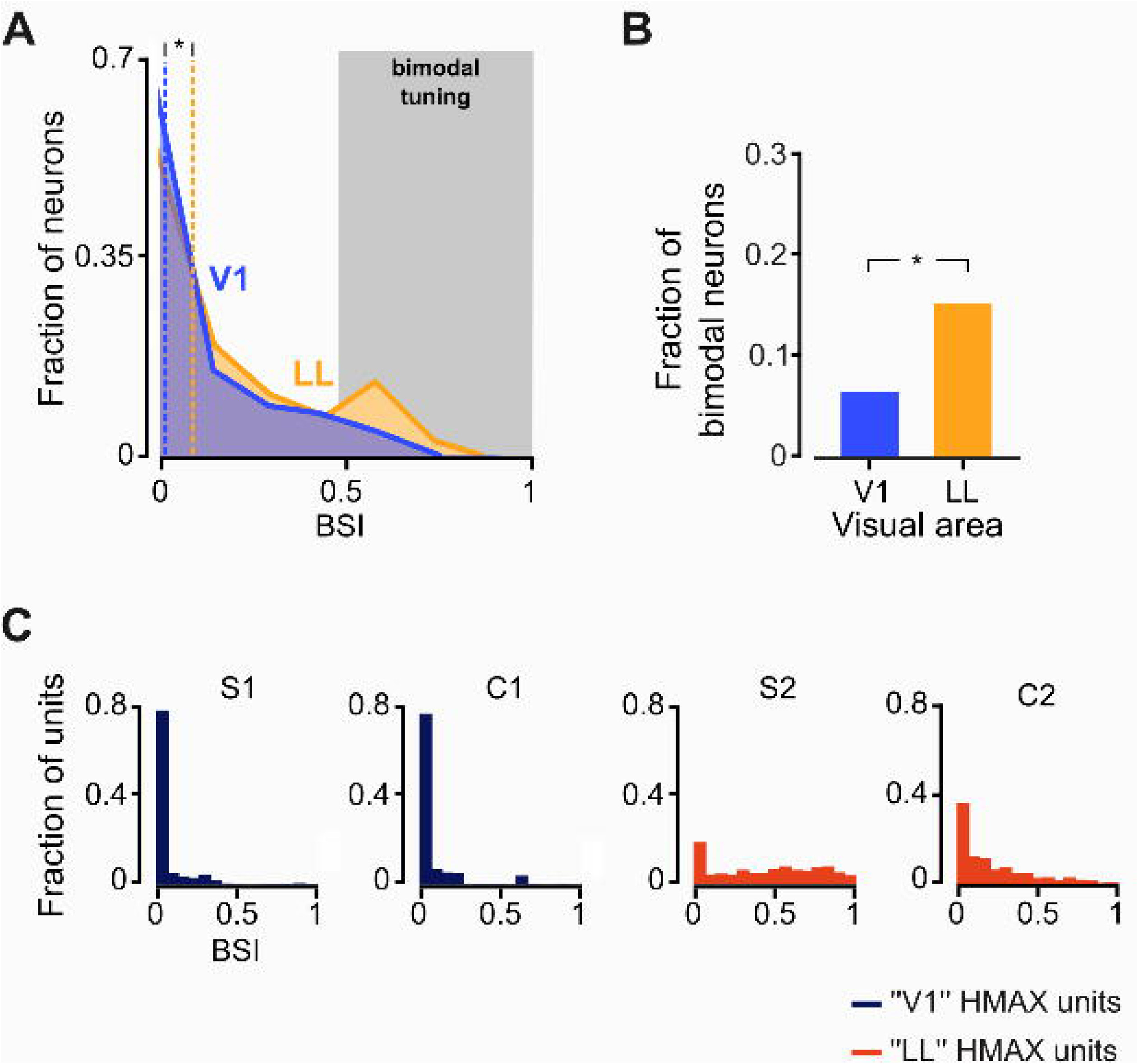
The entrainment of the neuronal response to the phase of drifting gratings decreases from V1 to LL. **A**. Distributions of the modulation index (MI) for the populations of V1 (blue; *n* = 105) and LL (orange; *n* = 104) neurons. Medians are shown as dashed lines (\*\*\**p* < 0.001; one-tailed Mann-Whitney U-test). The gray-shaded area highlights the region of the MI axis (MI > 3) corresponding to strongly phase-modulated units, i.e., simple-like cells. **B**. Fraction of simple-like cells (i.e., units with MI > 3) in V1 (blue) and LL (orange; \*\*\**p* < 0.001; *χ*^2^ test). **C**. Evolution of the MI distribution across the layers of the HMAX model (Fig. 3). Note the different scale on the ordinate axis for the S1 layer, as compared to downstream layers. **D**. Fraction of cells that have been reported as being strongly modulated by the phase of drifting gratings (i.e., simple-like cells) in areas V1 (blue; *n* = 10), V4 (brown; *n* = 3) and IT (orange; *n* = 1) of the monkey ventral stream (mean across *n* studies in each area ± SE; see Table 1). The drop of phase modulation across the three areas was statistically significant (*p* < 0.01, one-way ANOVA).

The agreement between the evolution of the modulation index observed in HMAX and in our data suggests the existence of specialized computations that build up transformation tolerance along rat lateral extrastriate cortex. To further support this conclusion, we checked whether the same qualitative trend has been reported in the monkey ventral stream literature. To this aim, we performed a simple meta-analysis of the evolution of phase-sensitivity to drifting gratings across areas V1, V4 and IT of primates. In V1, the fraction of simple cells reported by various authors is quite variable (Hubel and Wiesel, 1968; Schiller et al., 1976; De Valois et al., 1982a, 1982b; Foster K H et al., 1985; O’Keefe et al., 1998; Ringach et al., 2002; Kagan et al., 2002; Aronov et al., 2003; Gur et al., 2005), ranging from ~16% to 63% (Table 1), with a mean across studies of ~41% (blue bar in Fig. 4D). According to the few studies that investigated the phase-dependence of neuronal responses to grating stimuli in downstream areas (Table 1), the proportion of simple-like cells diminishes to ~14% in V4 (Desimone and Schein, 1987; Gallant et al., 1993, 1996), to become zero in IT (Pollen et al., 1984) (brown and orange bars in Fig. 4D), resulting in an overall significant decrement along the ventral stream (*p* = 0.0098, *F*_2,13_ = 7.25; one-way ANOVA). This trend, which is in agreement with the outcome of the HMAX simulations, confirms that the drop in the fraction of simple-like cells across consecutive visual processing stages can be taken as a strong marker of ventral stream computations. As such, its agreement with the trend observed from V1 to LL in our study (compare to Fig. 4B) suggests the existence of specialized machinery to build invariant representations of visual objects along rat lateral extrastriate cortex (see also Fig. 10A).

**Table 1.** Full list of research papers included in the meta-analysis shown in Figure 4D. The first column lists all the articles used in the meta-analysis by the name of the first authors and the publication dates. The second column indicates the cortical area that was studied in each individual article. The third column reports the neuronal sample size of each study. The fourth column shows the fraction of phase-modulated units reported in each article. The bars shown in Figure 4D were obtained by averaging the values in this column across studies within each individual cortical area. The fifth column specifies the criterion adopted in each study to define phase-modulated units (OI stands for Overlap Index, while F1/F0 is a classical measure of temporal modulation of the neuronal response to the frequency of drifting gratings, which is equivalent to the modulation index used in our study).

| **reference** | **area** | **number of units** | **fraction of modulated units** | **modulated units criterion** |
| --- | --- | --- | --- | --- |
| Hubel et al. 1968 | V1 | 272 | 0.45 | qualitative (overlap) |
| Schiller et al 1976 | V1 | 654 | 0.42 | qualitative (overlap) |
| De Valois et al. 1982a | V1 | 343 | 0.61 | $F1/F0 > 1$ |
| De Valois et al. 1982b | V1 | 222 | 0.63 | qualitative (drift modulation) |
| Foster 1984 | V1 | 122 | 0.28 | qualitative (overlap) |
| O'Keefe et al. 1998 | V1 | 210 | 0.46 | $F1/F0 > 1$ |
| Ringach et al 2002 | V1 | 308 | 0.40 | $F1/F0 > 1$ |
| Kagan et al 2002 | V1 | 114 | 0.22 | $OI < 0.3$ |
| Aronov et al. 2003 | V1 | 70 | 0.53 | $F1/F0 > 1$ |
| Gur et al 2005 | V1 | 339 | 0.16 | $OI < 0.3$ |
| DeSimone et al. 1987 | V4 | 322 | 0.16 | $F1/F0 > 1$ |
| Gallant et al 1993 | V4 | 28 | 0.11 | phase effect (ANOVA) |
| Gallant et al 1996 | V4 | 103 | 0.14 | phase effect (ANOVA) |
| Pollen et al 1984 | IT | 43 | 0.00 | qualitative (drift modulation) |

To further assess the involvement of nonlinear computations in establishing the tuning of LL neurons, we compared the quality of the RFs inferred through the STA method in V1 and LL. To this aim, each pixel intensity value in a STA image was z-scored, based on the null distribution of STA values obtained for that pixel, after randomly permuting the association between frames of the movie and spike times 50 times (Materials and Methods). This allowed expressing the intensity values of the STA image in terms of their difference (in units of standard deviation) from what expected in the case of no frame-related information carried by the spikes. We then computed a *contrast index* (CI; see Fig. 5A and a full description in Materials and Methods) to measure the amount of signal contained in the image and establish a conservative threshold (CI > 5.5), beyond which the linear RF structure of a neuron could be considered as well defined.

Figure 5C shows the 10 best STA-based RFs in each area, sorted according to the magnitude of their CI, from largest to smallest. In V1, most of these RFs featured equally prominent excitatory and inhibitory regions, often displaying a sharp, Gabor-like structure, while, in LL, the best STA images typically presented a single lobe, with much lower contrast. As a result, the median CI was higher for V1 than for LL neurons (blue vs. orange dashed line in Fig. 5D; *p* = 8.09*10^−10^; one-tailed Mann-Whitney U-test), and the fraction of units with well-defined linear RFs (i.e., with CI > 5.5) was five times larger in V1 than in LL (Fig. 5E; *p* = 6.48*10^−10^, χ^2^ test, *df=* 1).

**Figure 5.**
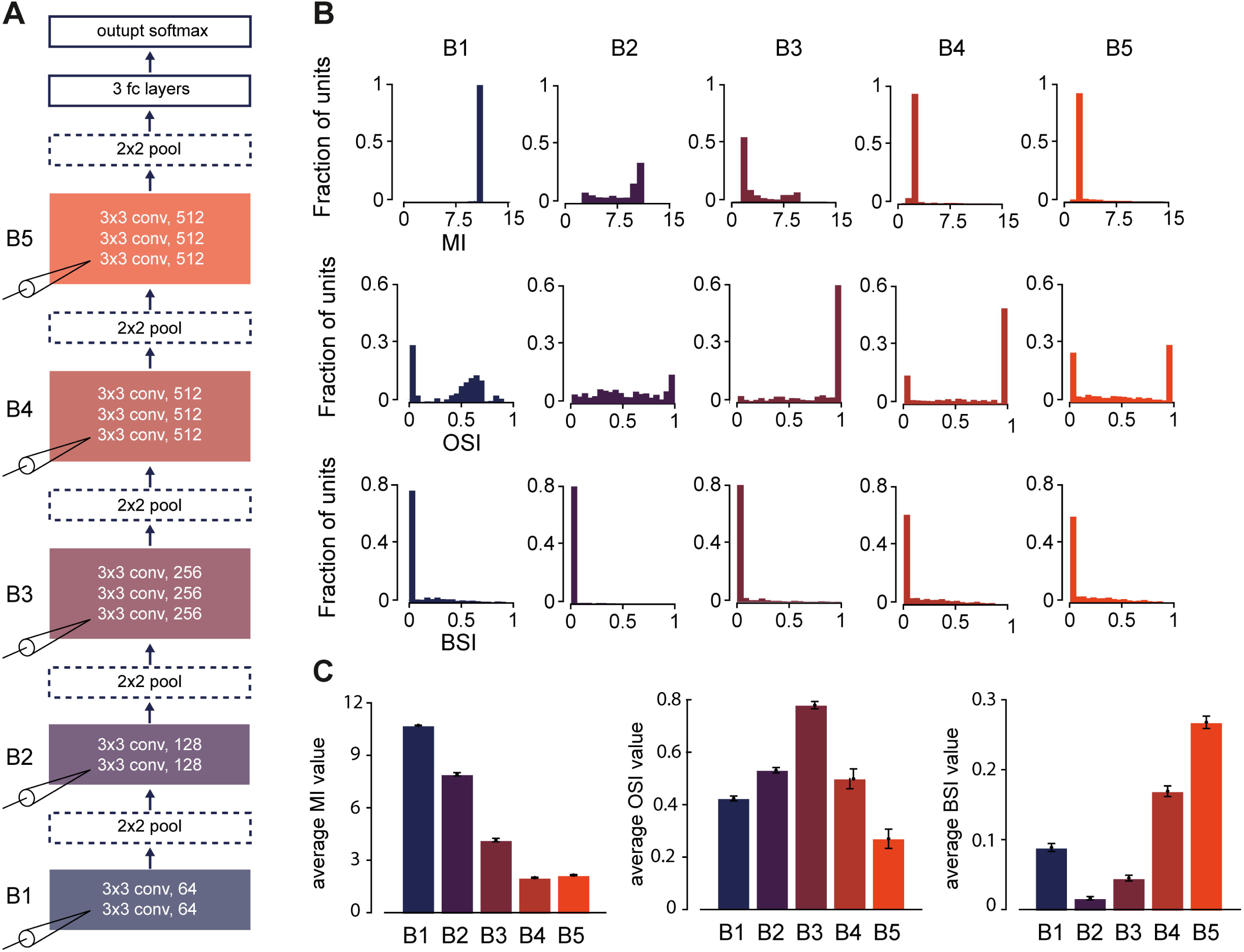
Spike-triggered average yields better-defined and structurally richer receptive fields in V1 than in LL. **A**. Illustration of the procedure to compute the contrast index of an STA image. The z-scored STA image obtained for a representative V1 neuron (left) is shown along the corresponding local-contrast map (middle), obtained by computing the contrast of the STA image (i.e., peak-to-peak difference) in local convolutional patches, with size equal to one fifth of the total image size (both images are cropped around the RF center). The contrast index (CI) associated to the STA image was computed by considering the resulting contrast distribution (right; bar plot) and taking its 0.9 quantile (dashed line; see Materials and Methods). **B**. Illustration of the procedure to count the lobes of an STA image. The output of the lobe counting algorithm is shown as a function of the binarization threshold (increasing from left to right) that was applied to the modulus of the z-scored STA image of the example neuron shown in **A**. Green dots show the centers of mass of the binarized images, whereas blue dots represent the centers of “valid” detected lobes (i.e., above-threshold, simply connected regions). Yellow dots mark “invalid” detected lobes (i.e., above-threshold, simply connected regions that were discarded for exceeding a distance limit from the center of mass of the binarized image; see Materials and Methods). For this example neuron, a very small spurious lobe was rejected at threshold = 3 (leftmost panel). **C**. Spatial structure of the ten best linear RFs obtained in V1 (top row) and LL (bottom row) using the STA method. The quality of each RF was assessed by computing the contrast index CI to quantify the amount of signal contained in a STA image (as illustrated in **A**). For each neuron, the RF shown here is taken at the time before the spike when CI was maximum (the corresponding CI value is reported above each STA image). As explained in Fig. 2, each pixel intensity value in a STA image was independently z-scored according to the null distribution yielded by a permutation test; the resulting z-scored values are reported here in units of standard deviation σ (gray scale bar). **D**. CI distributions for the populations of V1 (blue; *n* = 105) and LL (orange; *n* = 104) neurons. Medians are shown as dashed lines (\*\*\**p* < 0.001; one-tailed Mann-Whitney U-test). The gray-shaded area highlights the region of the CI axis (CI > 5.5) corresponding to units with well-defined linear RFs. **E**. Fraction of units with well-defined linear RFs (i.e., with CI > 5.5) in V1 (blue) and LL (orange; \*\*\**p* < 0.001; ***χ***^2^ test). **F**. Heat maps showing the distributions of lobe counts (abscissa) for the units with well-defined linear RFs (i.e., with CI > 5.5) in V1 (blue; *n* = 56) and LL (orange; *n* = 13) as a function of the binarization threshold (ordinate) used by the lobe-counting algorithm illustrated in B. Stars indicate threshold values for which the two distributions were significantly different (\**p* < 0.05, \*\**p* < 0.01; Fisher exact test).

We also observed a stark difference between the two areas in terms of the number of lobes contained in a STA image. To count the lobes, we binarized each image, by applying a threshold to the modulus of its intensity values (see Fig. 5B and a full description in Materials and Methods; also note that this analysis was restricted to the units with well-defined linear RFs). Regardless of the choice of the binarization threshold, the distribution of lobe counts peaked at 1 in both areas, but had a heavier tail in V1 than in LL (compare matching rows in Fig. 5F), resulting, in most cases, in a statistically significant difference between the two areas (Fisher’s exact test; * *p* < 0.05; ** *p* < 0.01; see legend for exact *p* values). Overall, this indicates that, in many cases, the responses of V1 neurons could be well approximated as a linear, weighted sum of the luminance intensity values falling inside their RFs – hence the sharp, multi-lobed STA images. By comparison, the lower-contrast, structurally simpler RFs yielded by STA in LL suggest a dominant role of non-linear computations in establishing the stimulus-response relationship in this area. In fact, in the case of a prevalently non-linear unit, the STA method would be able to reconstruct at most its linear sub-parts (Fournier et al., 2011, 2014; Sedigh-Sarvestani et al., 2014) – a sort of linear “residue” of the actual nonlinear filter. Such linear residue would still show, in some cases, the position and main orientation of the underlying filter, but would fall short at accounting for the full richness and complexity of its structure and for the extension of its invariance field (see next paragraph). This conclusion was supported by the tendency (although not significant) of the STA-based RFs to explain a lower fraction of variance of the orientation tuning curves in LL, as compared to V1 (Fig. 6).

**Figure 6.**
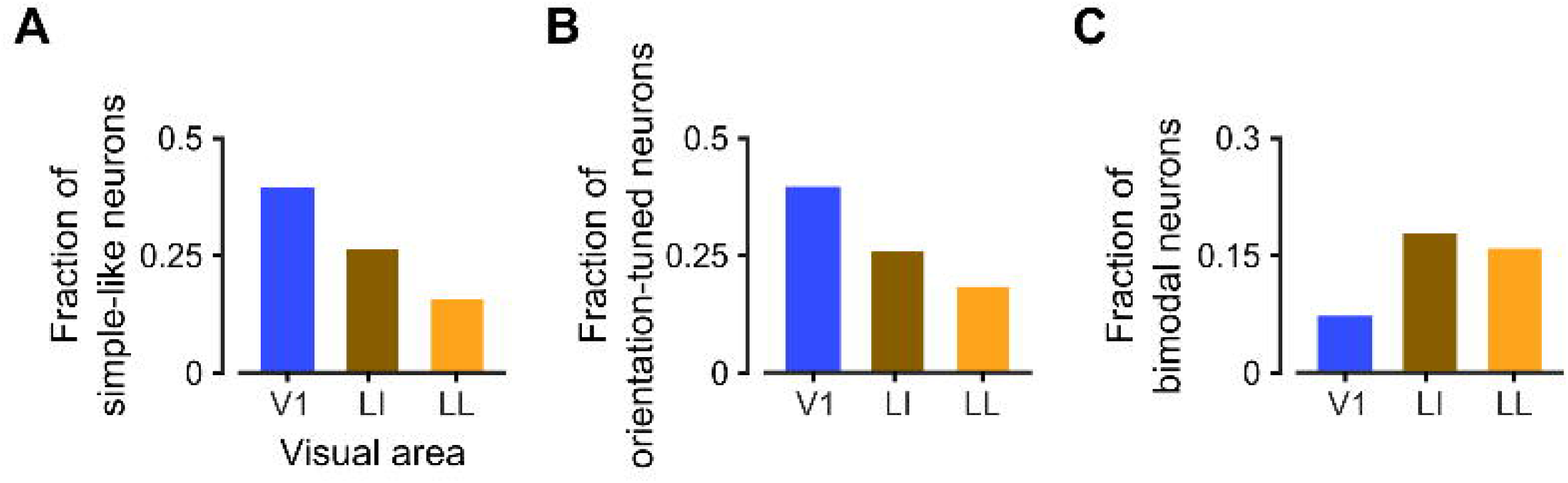
The V1 population includes neurons, whose direction-tuning curves are well predicted by linear LN models. **A**. Distributions of the variance of the direction-tuning curves that was explained by STA-based LN models for the populations of V1 (blue; *n* = 56) and LL (orange; *n* = 13) neurons with well-defined linear RFs (i.e., neurons with STA images having CI > 5.5; see Fig. 5D). Although the medians of the two distributions were not significantly different (*p* = 0.075; one-tailed Mann-Whitney U-test), a heavier tail can be noticed in the blue curve, which highlights the presence of a subset of units, in V1, whose selectivity was highly predictable using a linear, STA-estimated RF. **B**. Fraction of neurons, whose direction-tuning curves were well predicted by the STA-based LN models (i.e., units for which at least one third of the variance was explained by the models; gray patch in **A**) in V1 (blue) and LL (orange). Although twice as many well-predicted tuning curves were found in V1, as compared to LL, the difference was only marginally significant (*p* = 0.670; Fisher exact test).

The binarized STA images could also be used to compare the size (diameter) of the well-defined linear RFs in the two areas (with the diameter computed as the mean of the major and minor axes of the ellipse that best fitted the area covered by the detected lobes). When a liberal binarization threshold of 3.0 was chosen (with the goal of capturing most of the RF extension), the average diameter in LL (55.4 ± 10°; *n* = 13) was significantly larger than in V1 (39.1 ± 3.0°; *n* = 54; *p* = 0.02; one-tailed, unpaired t-test, *df* = 65). However, such difference became smaller (34.2 ± 2.9° in V1 vs. 39.4 ± 5.1° in LL) and not significant (*p* = 0.21; one-tailed, unpaired t-test, *df* = 65), as soon as the binarization threshold was raised at 3.5. This result should not be taken as evidence against the increase of RF size that is expected to take place along a ventral-like pathway. Such an increase has already been reported by several authors (Montero, 1981; Espinoza and Thomas, 1983; Vermaercke et al., 2014; Vinken et al., 2016) and was carefully quantified in our previous investigation of rat lateral extrastriate cortex (Tafazoli et al., 2017), where we found the RF diameter in LL (~30°) to be twice as large as in V1 (~15°). These estimates were obtained using a mapping procedure in which small, high-contrast stimuli (in our case, drifting bars with multiple orientations) were presented over a grid of visual field locations. While such high-contrast, localized stimuli can effectively elicit responses over the full invariance field of a neuron, thus yielding reliable RF estimates also for highly position-tolerant (i.e., nonlinear) units, the linear STA analysis used in our current study cannot, by definition, achieve this goal. In fact, for a highly tolerant neuron (such as a complex cell), a spike can be triggered both by a given distribution of luminance intensity values (a noise pattern) in a given portion of the visual field, and by its contrast reversal – which results in a failure of the STA method to map that portion of the RF. In other words, the nonlinear stimulus-response relationship that is typical of highly transformation-tolerant neurons prevents an accurate estimate of their RF size using STA. As a result, the RF size estimates yielded by STA in LL are largely underestimated, when compared to those obtained in V1, while the latter are consistent with the RF sizes that can be inferred from previous mouse studies, in which V1 receptive fields were also mapped using a linear reverse correlation method (Niell and Stryker, 2008; Bonin et al., 2011; Hoy and Niell, 2015).

To conclude, the poor quality of the STA-based RFs obtained in LL (in terms of contrast, number of lobes and estimated size; Fig. 5), taken together with the prevalence of complex-like units in this area (Fig. 4A-B), is consistent with the non-linear computations that are expected to build invariance along an object-processing hierarchy.

### Sharpness and complexity of orientation tuning follow opposite trends from V1 to LL

Next, we compared the V1 and LL neuronal populations in terms of the shape and sharpness of the orientation tuning curves obtained in the two areas. As shown in Figure 7A, in area LL most units had OSI lower than 0.5 (orange curve), with a median of 0.41 (dashed orange line), while in V1 the peak of the OSI distribution (blue curve) was in the 0.6-0.8 range, with a median of 0.53 (dashed blue line). The difference between the two OSI distributions was significant (*p* = 0.0029; Kolmogorov-Smirnov test), as it was the difference between their medians (*p* = 0.0084; one-tailed, Mann-Whitney U-test). Following a convention established in previous studies (Table 2) we classified as orientation-selective the units with OSI > 0.6, and we compared the fraction of units above such threshold in the two areas (Fig. 7B). In V1, the proportion of selective cells was twice as large as in LL (40% vs. 18%), and such difference was highly significant (*p* = 8.15*10^−4^; χ^2^ test, *df* = 1).

**Figure 7.**
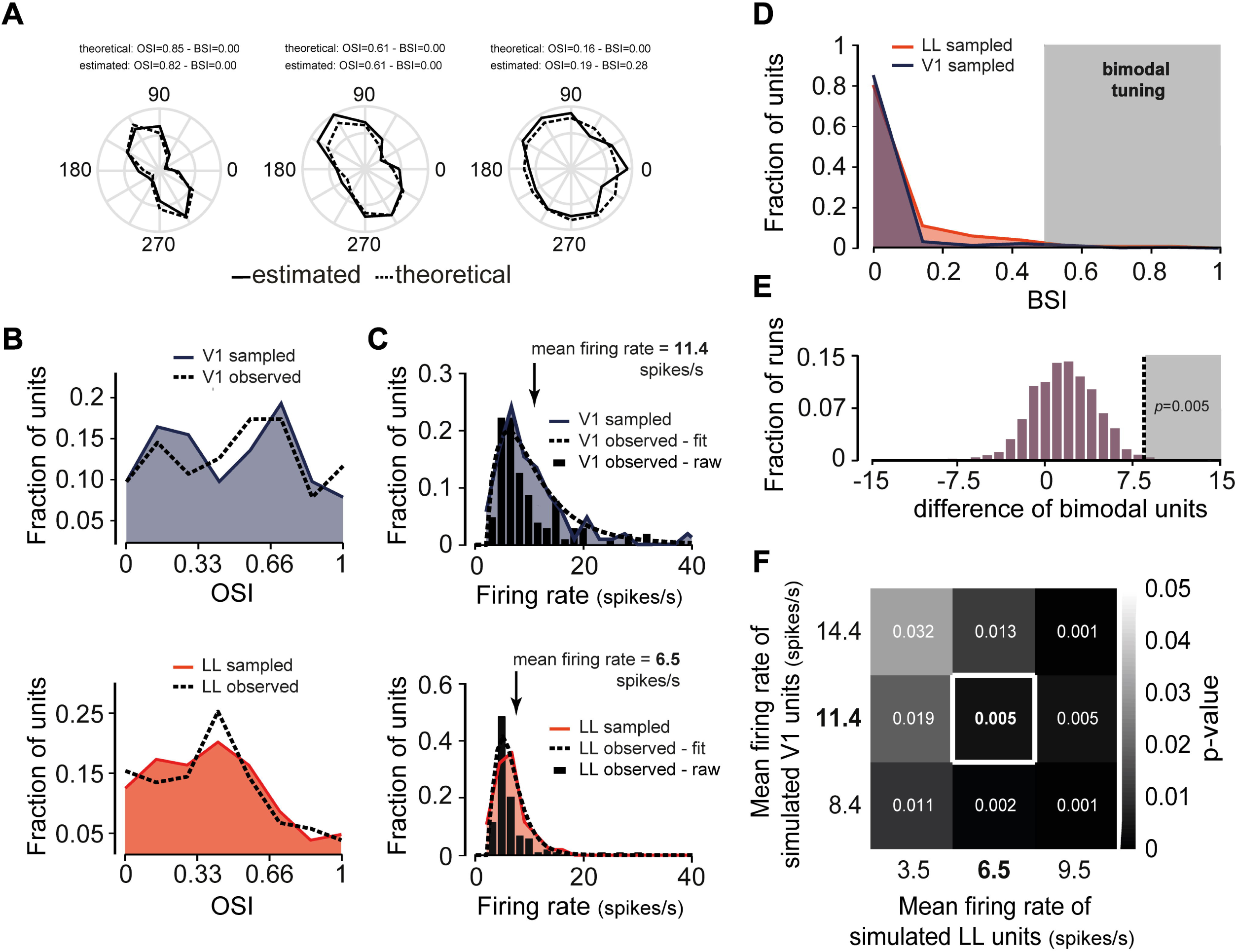
Orientation selectivity decreases from V1 to LL. **A**. Distributions of the orientation selectivity index (OSI) for the populations of V1 (blue; *n* = 105) and LL (orange; *n* = 104) neurons. Medians are shown as dashed lines (\*\**p* < 0.01; one-tailed Mann-Whitney U-test). The gray-shaded area highlights the region of the OSI axis (OSI > 0.6) corresponding to orientation-selective units. **B**. Fraction of orientation-selective cells (i.e., units with OSI > 0.6) in V1 (blue) and LL (orange; \*\*\**p* < 0.001; ***χ***^2^ test). **C**. Evolution of the OSI distribution across the layers of the HMAX model (Fig. 3). **D**. Fraction of cells that have been reported as being orientation tuned in areas V1 (blue; *n* = 7), V4 (brown; *n* = 4) and IT (orange; *n* = 4) of the monkey ventral stream (mean across *n* studies in each area ± SE; see Table 2). The drop of orientation tuning across the three areas was highly significant (*p* < 0.001, one-way ANOVA).

Although the computation of the OSI index is very popular in studies of neuronal tuning for oriented gratings (see Table 2), this metric, being based on the average responses of the neurons across repeated stimulus presentations, does not take into account the trial-by-trial variability of firing rates and, as such, does not provide a direct estimate of the information about orientation conveyed by neuronal firing. Therefore, we decided to complement the comparison of V1 and LL based on the OSI with an information theoretic analysis. For each neuron, we took its preferred direction (e.g., 60°) and the opposite one (e.g., 240°) and we assigned them to a stimulus category that we labeled as *preferred orientation*. We then took the two directions orthogonal to the preferred ones (e.g. 150° and 330°) and we assigned them to a category that we labeled as *orthogonal orientation*. We then pooled all the responses of the neuron to the stimuli within each category and we computed the mutual information *I*(*R;S*) between the neuronal response *R* (discretized into two equi-populated bins) and the stimulus category *S* (preferred vs. orthogonal). All quantities were computed using the Information Breakdown Toolbox (Magri et al., 2009) and were corrected for limited sampling bias using the Panzeri-Treves method (Panzeri and Treves, 1996; Panzeri et al., 2007). We found that V1 neurons conveyed, on average, more than twice as much information about orientation than LL units (i.e., 0.054 ± 0.007 bits vs. 0.023 ± 0.003 bits) and this difference was highly significant (*p* = 7.82^*^10^−5^; two-tailed, unpaired t-test, *df* = 207). Moreover, the amount of orientation information coded by the neurons within each population was positively and significantly correlated with OSI (*r* = 0.57, *p* = 2.48^*^10^−10^, *df* = 103 and *r* = 0.65, *p* = 4.52^*^10^−14^, *df =* 102 for, respectively, V1 and LL). Overall, these results confirm the validity of the analysis based on the OSI metric, and show that, even when the signal-to-noise ratio of neuronal responses is taken into account, V1 neurons display a much higher degree of orientation selectivity, as compared to LL units.

To check the consistency of this finding with ventral-like processing, we looked at the evolution of orientation tuning in HMAX and we carried out another meta-analysis of the monkey literature. As shown in Figure 7C, in the model, the OSI distribution underwent an abrupt shift towards much lower values in the transition from the C1 layer, corresponding to V1 complex cells, to the S2 layer, corresponding to a downstream area, where tuning for combinations of multiple oriented elements is achieved using a non-linear template matching operation (see Fig. 3). The shape of the OSI distribution was then maintained in the C2 layer, performing the max-pooling operation over the S2 afferents. With regard to the monkey ventral stream (Table 2), most authors report a very large fraction of orientation-tuned cells in V1 (Schiller et al., 1976; De Valois et al., 1982a, 1982b; Vogels and Orban, 1991; Zhou et al., 2000; Ringach et al., 2002; David et al., 2006), with a mean across studies close to 89% (blue bar in Fig. 7D). This proportion significantly decreases along downstream areas (*p* = 2.31^*^10^−6^, *F*_2_,_14_ = 46.17, one-way ANOVA), with ~65% and ~19% of orientation-tuned neurons reported, respectively, in V4 (Desimone and Schein, 1987; Pasupathy and Connor, 1999; Zhou et al., 2000; David et al., 2006) and IT (Desimone et al., 1984; Vogels and Orban, 1993, 1994; Sary et al., 1995) (brown and orange bars in Fig. 7D). These trends are confirmed by the few studies that applied the same experimental design to directly compare orientation tuning in V1 and either V4 (David et al., 2006) or IT (Vogels and Orban, 1991, 1993). Vogels and Orban found ~85% of orientation-tuned neurons in V1, compared to just ~14% in IT. David and colleagues reported a similar figure for V1, while in V4 they found that only 50% of neurons were orientation-tuned. In addition, they found that the fraction of cells with bimodal tuning (i.e., responding strongly to two non-adjacent orientations) was more than twice as large in V4 (28%) as in V1 (11%), in agreement with the increasing proportion of neurons that, in V4, are tuned for the combinations of oriented elements found in corners and curved boundaries (Gallant et al., 1993, 1996; Pasupathy and Connor, 1999, 2002).

**Table 2.** Full list of research papers included in the meta-analysis shown in Figure 7D. First three columns: same description as in Table 1. The fourth column shows the fraction of orientation-tuned units reported in each article. The bars shown in Figure 7D were obtained by averaging the values in this column across studies within each individual cortical area. The fifth column specifies the criterion adopted in each study to define orientation-selective units (BW stands for the bandwidth of the orientation tuning curve, while OSI is the same orientation selectivity index used in our study).

| **reference** | **area** | **number of units** | **fraction of tuned units** | **tuned units criterion** |
| --- | --- | --- | --- | --- |
| Schiller et al.1976 | V1 | 654 | **0.75** | BW < 90° |
| De Valois et al. 1982a | V1 | 343 | **0.90** | BW < 90° |
| De Valois et al. 1982b | V1 | 222 | **0.92** | BW < 90° |
| Vogels et al. 1991 | V1 | 285 | **0.85** | BW < 90° |
| Zhou et al. 2000 | V1 | 63 | **0.89** | OSI > 0.6 |
| Ringach et al. 2002 | V1 | 308 | **1.00** | BW < 90° |
| David et al. 2006 | V1 | 45 | **0.90** | BW < 90° |
| DeSimone et al. 1987 | V4 | 322 | **0.70** | BW < 90° |
| Pasupathy et al. 1999 | V4 | 152 | **0.52** | orientation effect (ANOVA) |
| Zhou et al 2000 | V4 | 45 | **0.82** | OSI > 0.6 |
| David et al. 2006 | V4 | 87 | **0.50** | BW < 90° |
| DeSimone et al. 1984 | IT | 66 | **0.00** | qualitative |
| Vogels et al. 1993 | IT | 958 | **0.13** | OSI > 0.7 |
| Vogels et al. 1994 | IT | 1885 | **0.34** | orientation effect (ANOVA) |
| Sary et al. 1995 | IT | 330 | **0.27** | orientation effect (ANOVA) |

Overall, the trends observed in our HMAX simulations (Fig. 7C) and emerging from the ventral stream literature (Fig. 7D) suggest that building tuning for complex visual patterns leads to a strong reduction of orientation tuning. As such, the drop of orientation selectivity observed from V1 to LL (Fig. 7A, B) can be taken as indication that LL neurons are tuned to more complex combinations of visual features, as compared to V1 units. Support to this conclusion came from comparing the two areas in terms of the tendency of the orientation tuning curves to be bimodal (Fig. 8A). In V1, most units had BSI equal or close to zero (i.e., virtually perfect unimodal tuning), with the distribution dropping sharply at larger BSI values (blue curve). In LL, the BSI distribution also peaked at zero, but featured a heavier tail, with a secondary peak above 0.5 (orange curve). As a result, the two distributions and their medians were significantly different (*p* = 0.044 and *p* = 0.029; Kolmogorov-Smirnov and one-tailed, Mann-Whitney U-test, respectively) and the fraction of neurons with bimodal tuning (i.e., with BSI > 0.5) was more than twice as large in LL as in V1 (*p* = 0.027; χ^2^ test, *df* = 1; Fig. 8B). Once again, a similar trend was observed in HMAX (Fig. 8C), where the BSI distribution displayed a much heavier tail in the deeper layers (S2 and C2) than in the early ones (S1 and C1).

**Figure 8.**
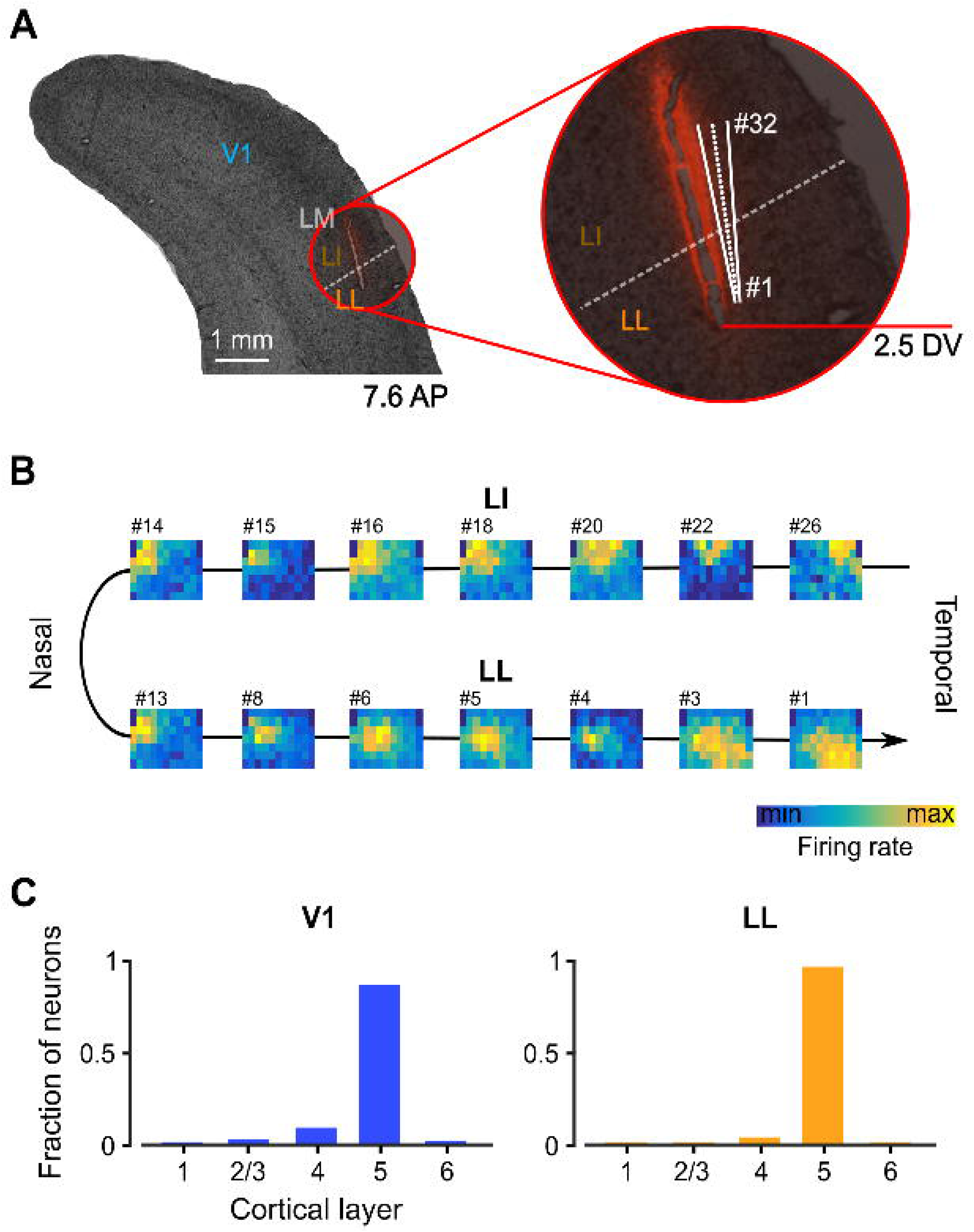
The tendency to be tuned for multiple orientations increases from V1 to LL. **A**. Distributions of the bimodal selectivity index (BSI) for the populations of V1 (blue; *n* = 105) and LL (orange; *n* = 104) neurons. Medians are shown as dashed lines (\**p* < 0.05; one-tailed Mann-Whitney U-test). The gray-shaded area highlights the region of the BSI axis (BSI > 0.5) corresponding to units with bimodal tuning. **B**. Fraction of cells with bimodal tuning (i.e., with BSI > 0.5) in V1 (blue) and LL (orange \**p* < 0.05; ***χ***^2^ test). **C**. Evolution of the BSI distribution across the layers of the HMAX model (Fig. 3).

Quantitatively, the difference in the number of bimodal neurons found in V1 and LL (7/105 vs. 16/104 in LL) may appear small, but, beside being statistically significant, is not dissimilar to the one reported in the monkey (David et al., 2006), when comparing V1 and V4 (see previous paragraph). In fact, while the incidence of bimodal neurons is approximately twice as large in monkey as in rat visual areas (11% vs. 7% in monkey vs. rat V1; 28% vs. 15% in V4 vs. LL), the percent increase from monkey V1 to V4 (~254%) is very similar to the percent increase from rat V1 to LL (~231%). In addition, in both V1 and LL, bimodal neurons came from multiple animals (4/12 in V1 and 4/6 in LL) and multiple recording sessions (4/15 in V1 and 5/9 in LL), which, not surprisingly (given with the low incidence of these neurons), accounted for a large fraction of all recorded, visually-driven single units in both areas (56/105 in V1 and 79/104 in LL).

Nevertheless, caution should be taken when comparing the BSI distributions obtained for two neuronal populations with a different level of orientation selectivity, because the BSI and OSI indexes are not constrained to be statistically independent. Intuitively, in presence of some noise, poorly selective neurons (i.e., cells with low OSI) will tend to have slightly larger BSI values than sharply selective units, even if the underlying tuning curves are unimodal (i.e., even if they are ideal oriented-edge detectors without any true secondary peaks in their orientation tuning curve). This is because, for a very broad tuning curve, the two largest, non-adjacent peaks will tend to be closer, in terms of magnitude, than for a narrow tuning curve. In our data, BSI and OSI were in fact negatively correlated (*r* = −0.48 and *r* = −0.4, respectively, in V1 and LL; *p* = 2.3^*^10^−7^, *df* = 103 and *p* = 1.1^*^10^−5^, *df* = 102; unpaired, two-tailed, t-test,). Therefore, we performed an analysis to quantitatively check that the larger BSI values found in LL, as compared to V1 (Fig. 8A), were not simply a byproduct of the lower orientation tuning of the LL neurons (Fig. 7A). In other words, we verified that the larger fraction of units being classified as bimodal in LL, as compared to V1 (Fig. 8B), was not an artifact of the possibly larger proportion of LL neurons with very broad (but unimodal) orientation tuning.

Our analysis was based on simulating two artificial populations of perfectly unimodal units (see examples in Fig. 9A), having: 1) the same size of the recorded V1 and LL populations (i.e., 105 and 104 simulated V1 and LL units, respectively); 2) OSI values sampled from the empirical OSI distributions that were observed in the recorded populations (dashed lines in Fig. 9B; same as the distributions shown in Fig. 7A); and 3) peak firing rates (FRs) sampled from log-normal functions (dashed lines in Fig. 9C) that were fitted to the empirical distributions of peak firing rates observed in the recorded populations (black bars in Fig. 9C). As a consequence of point 3 above, the average FRs of the two simulated populations were very close to those of the recorded populations (i.e., 11.4 ± 10.7 spikes/s in V1 and 6.5 ± 4.3 spikes/s in LL; see arrows in Fig. 9C). In addition, to account for the effect of estimating orientation-tuning curves from a limited number of repeated trials in noisy neurons, the responses of a simulated unit to a given orientation were drawn from a Poisson distribution with the mean equal to the theoretical value of the simulated tuning curve at that orientation (dashed lines in Fig. 9A). Ten repeated Poisson responses were simulated for each orientation, and then averaged to obtain sampled tuning curves for the simulated units (solid lines in Fig. 9A). The goal of our simulation was to check if, as a consequence of the Poisson noise, these sampled tuning curves would develop spurious secondary peaks (especially in the case of low orientation tuning) that could lead to artifactual differences of bimodal neurons between LL and V1. It should be noticed that simulating noisy Poisson neurons, along with the fact that a limited number of units were drawn from the theoretical, data-matched OSI and FR distributions, brought the sampled OSI and FR distributions to be slightly different from the theoretical ones. Such differences, however, were minimal, as it can be appreciated by comparing the sampled and theoretical OSI and FR distributions obtained for two example simulated V1 and LL populations in Figure 9B and C (colored lines/areas vs. black dashed lines). Figure 9D reports instead the sampled BSI distributions that were obtained for these example simulated populations. As expected (see previous paragraph), the larger orientation tuning of the simulated LL units did bring their BSI distribution (red curve) to have a slightly heavier tail, as compared to the BSI distribution of the simulated V1 units (blue curve). However, the difference between the two distributions was very minor, as compared to the one observed for the actual populations (compare to Fig. 8A). Most noticeably, the simulated LL distribution lacked the heavy tail, with the secondary peak at BSI > 0.5, that we found for the recorded LL neurons. As a result, the number of units classified as bimodal in these two simulated populations (i.e., with BSI > 0.5) was identical (3 units).

Given the stochastic nature of our analysis, this outcome does not guarantee that, in general, no major differences can be observed in the number of bimodal neurons found in the two simulated populations. Therefore, to test whether we could statistically reject the null hypothesis that the difference of bimodal units observed between the recorded LL and V1 populations was merely due to differences in terms of broadness of orientation tuning and firing rate magnitude (and, as such, response noisiness), we repeated the simulation illustrated above in 1,000 independent runs, so as to obtain 1,000 comparisons among BSI distributions of simulated LL and V1 units. In each run, we computed the difference between the numbers of neurons classified as bimodal in the two simulated populations. This yielded a null distribution of differences of bimodal units between LL and V1, under the null hypothesis of purely unimodal tuning in both areas (purple bars in Fig. 9E). When we compared the actual, measured difference of bimodal neurons recorded in LL and V1 (dashed line in Fig. 9E) to this null distribution, we found that the probability of getting such a difference (or a larger one) under the null hypothesis was *p* = 0.005. Therefore, we could statistically reject the hypothesis that the larger fraction of bimodal neurons observed in LL was only due to the lower level of orientation tuning observed in this area, as compared to V1, with a 0.005 significance level. Interestingly, this conclusion held true, even when we simulated LL and V1 populations with the same OSI distributions observed in our recordings, but with mean peak firing rates either 3 spikes/s smaller or 3 spikes/s larger than those measured in the two areas (Fig. 9F). In particular, we found that the difference of bimodal neurons between LL and V1 was significantly larger than expected by chance, also when the statistical comparison was performed against a null distribution obtained from LL and V1 simulated units with peak firing, respectively, lower and larger than that observed in the recorded populations (*p* = 0.032; top-left cell in Fig. 9F). That is, not matter how noisy the tuning curves of the simulated LL units were made with respect to those of the simulated V1 units, their tendency to display spurious secondary peaks was never so strong as to explain the difference of bimodal units observed in our recordings. Overall, this indicates that the majority of neurons classified as bimodal in LL were truly so, thus confirming that the different proportion of bimodal cells in LL and V1 (Fig. 8B) is a small, but robust effect.

**Figure 9.**
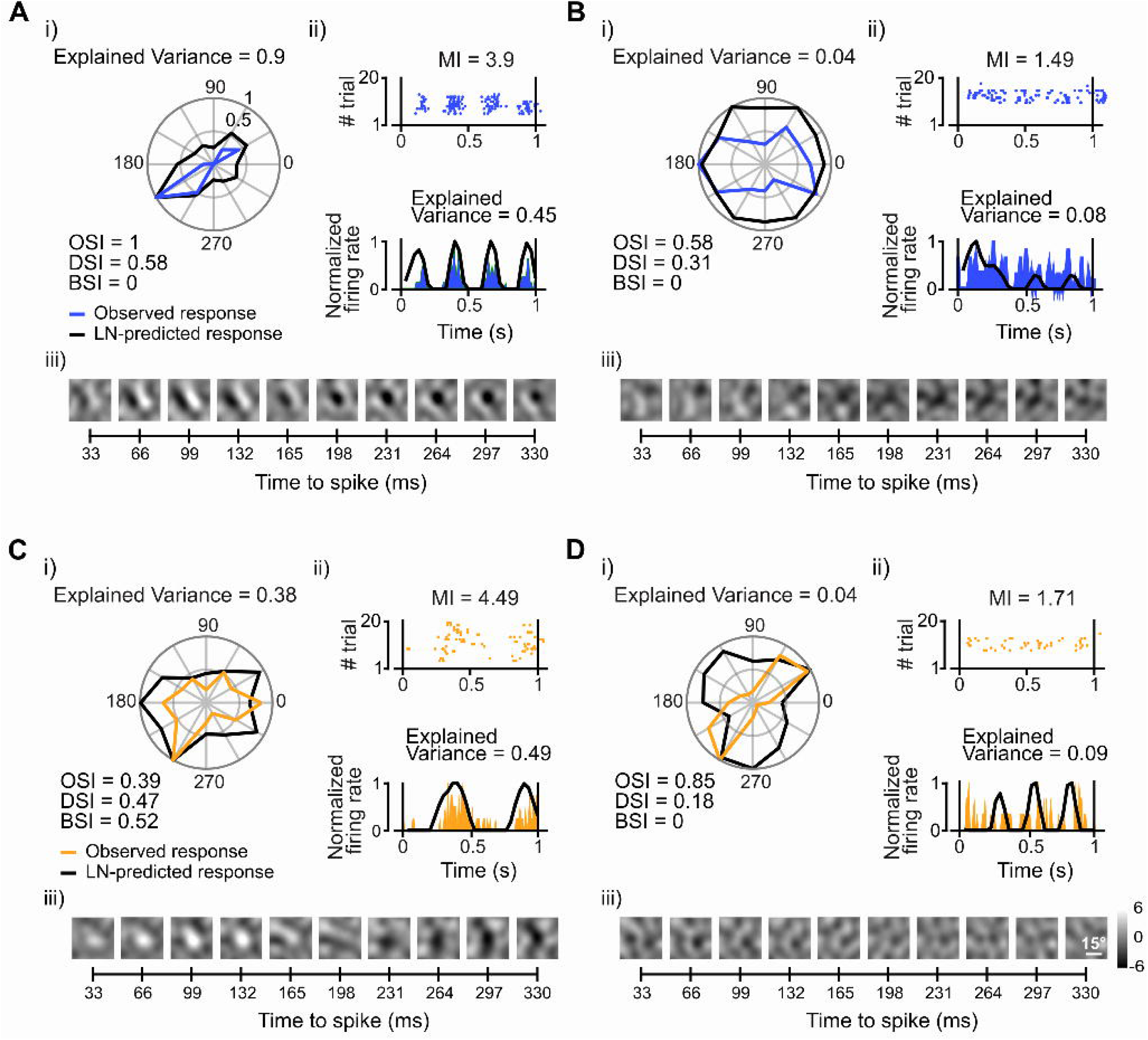
The increase of neurons with bimodal orientation tuning from V1 to LL is not accounted for by the broader orientation selectivity and lower firing rate of LL units. **A**. Examples of simulated units with unimodal orientation tuning curves, having a different degree of sharpness. Each tuning curve was obtained by summing two Von Mises functions (Kohn and Movshon, 2004), centered on the two opposite directions that were collinear with a given chosen orientation. The width of the Von Mises functions, which are the circular analogues of Gaussian functions, is controlled by the parameter *k*, which, in turn, is equivalent to the inverse of the variance of a Gaussian distribution (Berens, 2009). Starting from these theoretical tuning curves (dashed lines), we sampled 10 repeated Poisson responses for each orientation and we averaged these responses to obtain estimated tuning curves for the simulated units (solid lines). From these estimated tuning curves we computed the BSI of the simulated units. **B**. The OSI of each simulated unit was sampled from theoretical distributions that were identical to those obtained for the recorded V1 and LL populations (black dashed lines; same as the distributions shown in Fig. 7A). Obviously, as a result of the Poisson noise in the simulated responses (see **A**), the sampled OSI distributions obtained in any given run of the simulation (colored curves) were slightly different from the theoretical ones (the colored lines/areas show such sampled distributions for two example simulated populations). **C**. The peak firing rates (FRs) of the simulated units were sampled from lognormal functions (black dashed lines) that were fitted to the FR distributions obtained for the recorded V1 and LL populations (black bars). Again, because of the Poisson noise, the sampled FR distributions obtained in any given run of the simulation were slightly different from the theoretical ones (the colored lines/areas show such sampled distributions for same simulated populations shown in **B**). **D**. Sampled BSI distributions obtained for the same two example populations of simulated V1 (105) and LL (104) units shown in **B** and **C**. Despite the tendency of the simulated LL units to have slightly larger BSI values than the simulated V1 units, the fraction of units classified as bimodal (i.e., with BSI > 0.5) was the same (3 units) for these example populations. **E**. Simulated BSI distributions as those shown in **D** were obtained in 1,000 independent runs, thus yielding a null distribution of differences between the number of neurons classified as bimodal in the two populations (purple bars), under the hypothesis of purely unimodal tuning curves. Such null distribution was used to assess the significance of the measured difference of bimodal neurons between LL and V1 (*p* = 0.005; dashed line). **F**. The statistical test described in **E** was repeated for six different combinations of mean peak firing rates in the simulated V1 and LL populations. These six combinations included the mean FR values actually observed in the recorded populations (central square; same analysis shown in **E**). The additional combinations were obtained by either lowering or increasing the mean FR observed in V1 and LL by 3 spikes/s. For every combination, the difference of bimodal units between the two areas was significantly larger than what expected by chance (*p* < 0.05), under the null hypothesis of purely unimodal tuning curves (*p* values reported in the cells of the matrix; see also the color bar).

Interestingly, the relative small incidence of bimodal units in high-order areas is fully consistent with the HMAX simulations, where these units were ~51% in layer S2, but reduced to ~20% in layer C2 (Fig. 8C). This indicates that the max-pooling operation that builds invariance (yielding the C2 layer) partially counterbalances the template-matching computation that builds tuning for multiple features (yielding the S2 layer), when it comes to produce bimodal tuning. Taken together, these considerations support the interpretation that the decrease of OSI from V1 to LL, along with the concomitant increase of BSI, results from an increment of the complexity of tuning of LL neurons – a property that is consistent with their involvement in ventral processing.

This conclusion was confirmed by the overall lower effectiveness of the grating stimuli to drive LL neurons, as compared to V1 units (see Fig. 9C). By contrast, only a minor difference was observed in terms of response latencies (102 ± 11 ms in V1 vs 118 ± 9 ms in LL; see Materials and Methods). This is in disagreement with previous studies reporting lower latencies in both areas and larger differences between V1 (~40 ms) and LL (~75 ms) (Vermaercke et al., 2014; Tafazoli et al., 2017), as expected along a partially feedforward processing hierarchy. Such discrepancy is explained by the fact that full-field drifting gratings, as opposed to the localized, high-contrast visual shapes used in (Vermaercke et al., 2014; Tafazoli et al., 2017), are not suitable to properly estimate response latencies. In fact, as shown by the example neurons of Figure 2, the onset of the neuronal response is determined by time of the drift cycle in which the grating happens to properly align with the RF of the neuron. This, in turn, depends on the initial phase of the grating relative to the neuron’s RF (which varied randomly from neuron to neuron in our experiment, being fixed relative to the stimulus display) and on its spatial and temporal frequencies. As a result, response latencies measured with drifting gratings are highly variable – they largely overestimate the actual time at which neurons in a given area start processing visual inputs, and do not allow appreciating differences among areas.

Finally, we also measured the direction selectivity of the two neuronal populations, finding no significant differences in terms of DSI (0.42 in V1 vs. 0.45 in LL; *p* = 0.7; Mann-Whitney U-test), which suggests a lack of specialization of LL for motion processing, as compared to V1.

### Laterointermediate visual area (LI) displays a level of specialization for ventral processing that is intermediate between V1 and LL

Our recordings were aimed at area LL, since, in our previous study, this region bore the clearest signature of ventral processing within rat lateral extrastriate cortex (Tafazoli et al., 2017) – i.e., the lowest sensitivity to low-level image attributes (such as luminance and contrast) and the most invariant coding of object identity. A similar advantage, over V1, to support invariant object recognition was also displayed, although to a lesser extent, by the laterointermediate visual area (LI), which borders LL medially. Our current study was not designed to obtain a statistical characterization of LI in terms of tuning for drifting gratings. Nevertheless, we recorded a small number of single units also from this region, since, while aiming at LL, a fraction of the recording sites of the silicon probe landed also in LI (see Fig. 1). We therefore looked at the sensitivity for the phase of drifting gratings and at the sharpness and shape of the orientation-tuning curves also for the recorded LI neurons (a total of 23 responsive and reproducibly driven units).

**Figure 10.**
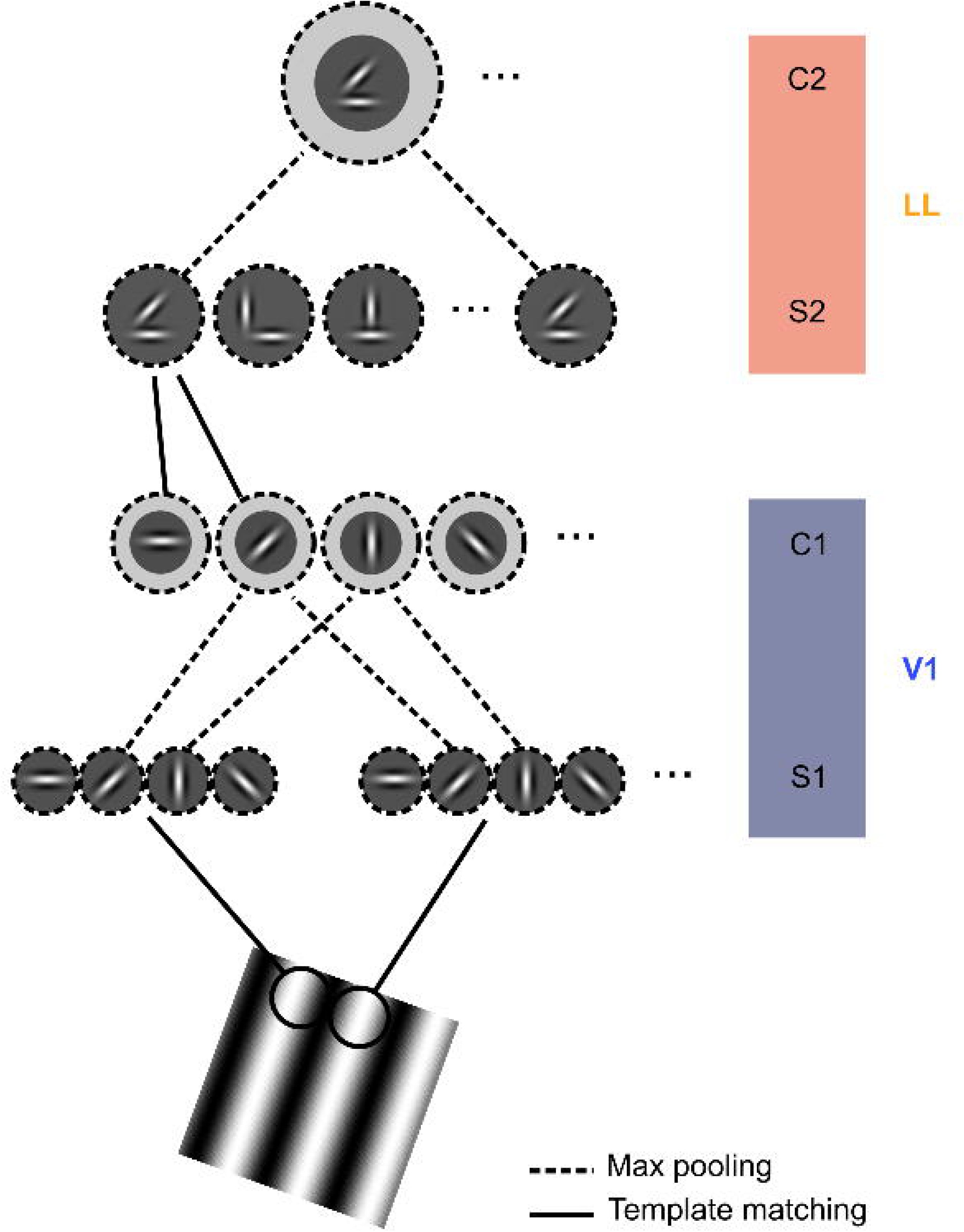
The level of specialization for ventral processing of LI is intermediate between V1 and LL. The fraction of LI neurons (brown bar) with simple-like behavior (left), sharp orientation tuning (middle) and bimodal tuning (right) is shown along the fractions of units displaying such properties in V1 and LL (blue and orange bars, respectively; same bars already shown Figs. 4B, 7B and 8B). No statistical comparison was performed, given the low sample of LI units available (23 responsive and reproducibly driven units).

As shown in Figure 10 (leftmost panel), the fraction of phase-modulated, simple-like neurons in LI (brown bar) was intermediate between V1 and LL. The same applied to the fraction of orientation-selective units (middle panel), while the proportion of neurons with bimodal tuning (rightmost panel) was similar in LI and LL. We did not quantify statistically these trends, given the low number of cells sampled from LI. Nevertheless, we report them here, because they are consistent with our earlier finding that the processing leading to the specialization of LL for ventral computations takes place gradually, along a processing chain involving multiple areas, of which LI is one of the nodes (Tafazoli et al., 2017).

### Evolution of the tuning for drifting gratings in a state-of-the-art deep convolutional network

As shown in the previous sections, most trends of variation observed along rat lateral extrastriate areas, in terms of tuning for drifting gratings, are in qualitative agreement with the prediction of HMAX. Such agreement is not surprising, given the selectivity-building and tolerance-building computations implemented by the units of the model. At the same time, the trends observed in HMAX are not trivial and should still be considered as emergent properties of the model, because, when such computations are implemented separately by thousands of independent units across a cascade of feedforward layers, the outcome of their interaction is not easily predictable. For instance, it is far from obvious that the template-matching computation that increases the complexity of tuning of S2 units will still allow them to show some selectivity for oriented gratings (Fig. 7C), and that such selectivity (also in layer C2) will be as phase-invariant as the one found (by construction) in layer C1 (Fig. 4C). Also, it was interesting to notice how the incidence of bimodal units was boosted (as expected) by the selectivity-building computation in layer S2, but it was dampened by the tolerance-building pooling performed by C2 units (Fig. 8C), so that, in this layer, the fraction of bimodal units was not far from the one we observed in LL (see previous section).

HMAX, however, is only one of the possible feedfoward neuronal networks with a brain-inspired architecture that can serve as a comparison for physiology/behavioral experiments. In particular, HMAX can be considered as one of the ancestors of modern deep convolutional neuronal networks (DCNNs), which, in recent years, have revolutionized the field of machine learning, matching (and even surpassing) human accuracy in a number of object recognition tasks (LeCun et al., 2015). DCNNs share the same basic architecture of HMAX, with alternating layers of complexification of feature selectivity and tolerance-increasing pooling, but they are much larger, being made of tens of layers, each with hundreds of thousands units (for a total of several millions units). More importantly, while HMAX has a static, hard-wired connectivity (Materials and Methods), the units in DCNNs iteratively adjust their synaptic weights, in such a way to learn feature representations that maximize their accuracy in a given image classification task (LeCun et al., 2015). Therefore, the emergent properties of DCNNs (e.g., in terms of shape selectivity) are less determined by architectural/computational constraints, as compared to HMAX, and more strongly driven by the optimization process underlying the development of powerful representations for object recognition. This has led several neuroscientists to use such networks as benchmarks to study the emergence of selective, yet invariant, representations of visual objects in the ventral stream of both monkeys (Yamins et al., 2014; Yamins and DiCarlo, 2016; Hong et al., 2016) and humans (Khaligh-Razavi and Kriegeskorte, 2014; Kriegeskorte, 2015; Kubilius et al., 2016; Kheradpisheh et al., 2016). Inspired by these primate studies, we repeated the same analysis carried out in HMAX also in VGG16, a state-of-the-art DCNN that scored second place in object classification and first place in object localization at the ILSVRC-2014 competition (Simonyan and Zisserman, 2014).

VGG16 is a quite large DCNN, totaling about 138 million parameters across its 16 convolutional and fully-connected layers (Fig. 11A). These layers are grouped in 5 *blocks* (named B1 to B5 in Fig. 11A), each composed by 2 or 3 convolutional layers (colored boxes) and followed by a max pooling layer (white boxes), with a final stack of 3 fully-connected layers on top, followed by a 1000-way softmax output layer for classification. Thanks to this relatively simple and uniform structure, VGG16 is particularly appealing as a model of how modern DCNNs work.

**Figure 11.**
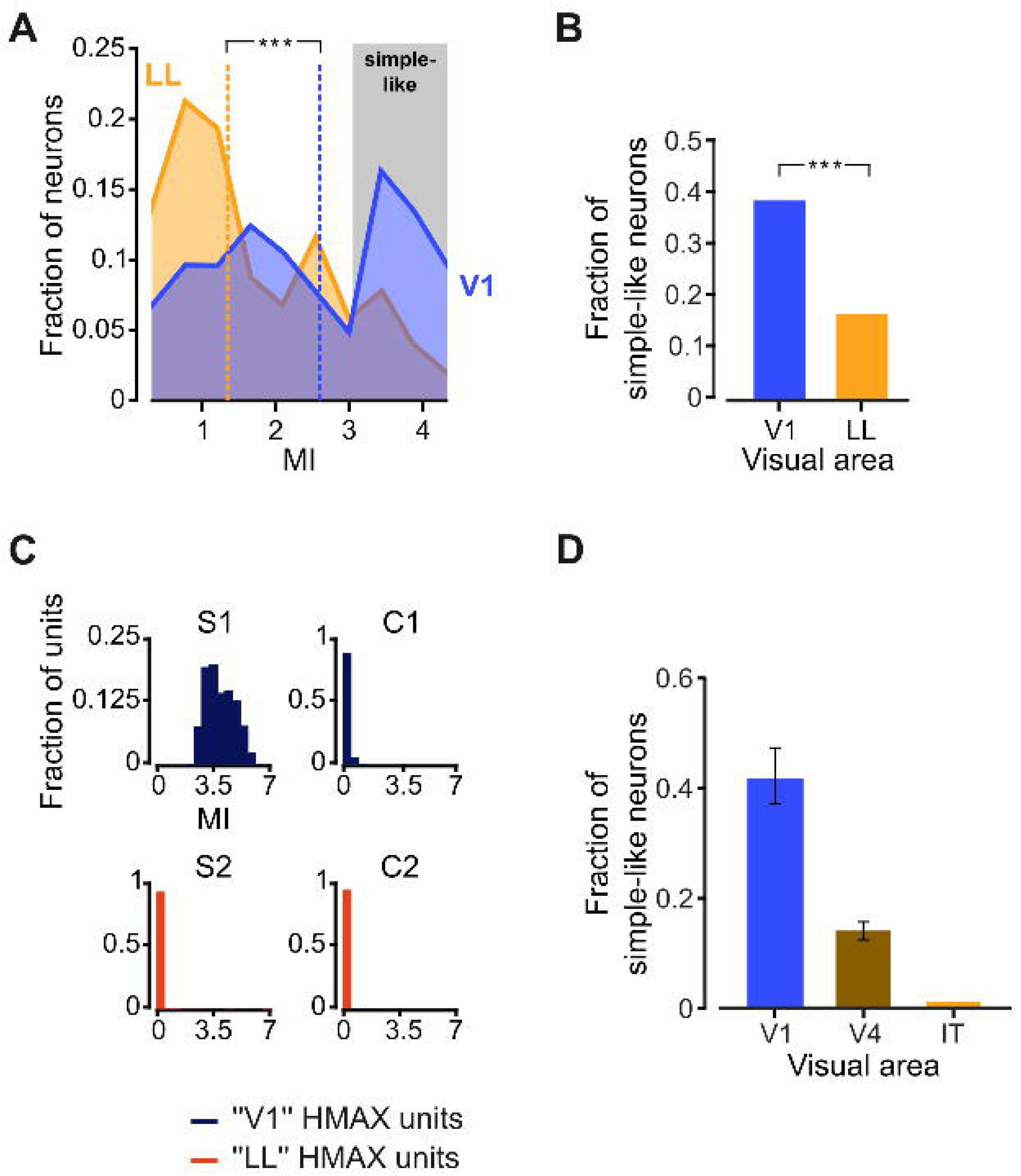
Evolution of the tuning for drifting gratings across the layers of a deep convolutional network. **A**. Sketch of the architecture of VGG16, a feedforward, deep convolutional network made of five blocks of 2 or 3 convolutional layers, followed by a max pooling layer, with a stack of three additional, fully-connected layers on top, before the softmax output layer for classification. In the drawing, the five blocks of convolutional layers (named B1 to B5) are color coded (from blue to red) to emphasize the progression from lower-(blue) to higher-order (red) representations of visual features implemented by the network. Each block also reports the size and number of the convolutional filters contained in each layer. Similar figures are reported for the pooling layers and for the upper, fully-connected layers (white frames). The weights of the network were those originally obtained by training it to correctly classify the object categories of a large image dataset (ImageNet). In our tests, the input layer of the network was fed with drifting gratings spanning different orientations, spatial frequencies and temporal frequencies. The tuning for key processing properties (MI, OSI and BSI) was then measured for a sample of 1000 randomly selected units from the first convolutional layer of each block (as highlighted by the picture of a probe shown over each sampled layer). **B**. Distribution of the MI (top), OSI (middle) and BSI (bottom) indexes across the five selected convolutional layers of VGG16 (same color code as in A). **C**. Average values (mean ± SE) of the MI (left), OSI (middle) and BSI (right) indexes over the pools of 1000 units sampled from of each of the five selected convolutional layers of the network (same color code as in A).

For our tests, we used VGG16 pre-trained on the ImageNet dataset (a large photographic database very popular as a computer vision benchmark), and, as done with HMAX, we fed to the input layer of the network a set of drifting gratings spanning the same range of directions used in the neurophysiology experiment (Materials and Methods). We then computed the activation (over the entire duration of each presented grating) of a pool of 1000 randomly selected units from the first convolutional layer of each block (Fig. 11A). For each unit, we measured the time evolution of its activation, so as to estimate its sensitivity to the phase of the drifting gratings through the MI index, and we obtained orientation-tuning curves, from which we computed the OSI and BSI indexes.

Figure 11B shows how the distributions of these indexes evolved across the five sampled layers of the network. In the case of the modulation index (top row), we observed a progressive shift from large (> 7.0) to low (< 3.0) MI values, resulting in a monotonic decrease in the mean level of phase modulation along the network (Fig. 11C, leftmost panel). This trend was very similar to the one displayed by HMAX (Fig. 4C), and was consistent with the drop of simple-like cells observed along the primate ventral stream (Fig. 4D) and, in our experiments, from rat V1 to LL (Fig. 4A-B and Fig. 10, leftmost panel). This confirms that the reduction of phase-modulated units can be taken as a strong marker of the existence of nonlinear, tolerance-building computations along rat lateral extrastriate cortex, similar to the max pooling implemented both in HMAX and VGG16.

The orientation selectivity index displayed instead a non-monotonic trend (Fig. 11B, middle row and Fig. 11C, middle panel), since it first increased, reaching a peak in B3, the third block of convolutional layers, and then decreased from B3 to B5. This behavior is qualitatively different from that observed in HMAX (Fig. 7C) and may seem inconsistent with the monotonic decrease of OSI reported across the monkey ventral stream (Fig. 7D) and observed, in our study, along rat lateral extrastriate cortex (Fig. 7A-B and Fig. 10, middle panel). Such discrepancy, however, is easily explainable by the fact that in VGG16, differently from HMAX, the first processing layer does not contain sharply-tuned oriented Gabors (of tens by tens pixels in size), but a first set of very small convolutional filters (of only 3x3 pixels). Having, by construction, such a small receptive field size, these units cannot learn the Gabor-like filters with high aspect ratio that are necessary to achieve sharp orientation tuning. Only the units in the following convolutional layers, as the effective receptive field size increases, can gradually evolve into sharply tuned, oriented-edge detectors. In other words, while HMAX, by design, is meant to simulate the processing that takes place from primary visual cortex onward, VGG16 could be regarded as a model of the entire visual system, with the initial layers, operating on the raw pixel representation of the image, being architecturally and functionally closer to subcortical visual areas (such as retina and LGN) than to the cortex.

In the light of these considerations, it is tempting to interpret the initial increase of OSI in VGG16 as the attempt of the network to first learn a bank of oriented-tuned units. Once this V1-like representation is established (in B3), the network displays a behavior very similar to HMAX and to monkey (rat) temporal (lateral) visual cortical areas, with OSI that gradually and monotonically decreases, as multiple peaks start appearing in the orientation tuning curves – a sign that the units of the upper layers are learning more complex feature representations. This was confirmed by the trend observed for BSI (Fig. 11B, bottom row and Fig. 11C, rightmost panel), which initially decreased (from B1 to B2), but then monotonically increased, with the fraction of units with bimodal tuning reaching ~25% in B5, a value not dissimilar from the one found in HMAX (se previous section). It is possible that DCNNs based on slightly different architectures, such as Alexnet (Krizhevsky et al., 2012), where the initial layer contains larger convolutional filters, may display a behavior that is even more consistent with HMAX, and show a strictly monotonic decrease of OSI and increase of BSI across consecutive layers (we did not test this possibility, since a systematic comparison of different DCNN architectures is clearly beyond the scope of our study).

Overall, we can conclude that all the trends we observed along rat lateral extrastriate cortex are in qualitative agreement not only with the predictions of the conceptual, hard-wired model of ventral processing implemented in HMAX, but also with the behavior displayed by the upper layers of a state-of-the-art, deep convolutional network trained for image classification. This adds further support to our conclusion that rat lateral visual cortical areas are specialized for processing object information.

## Discussion

In this study, we sought to validate and further explore the specialization of rat area LL for ventral computations, as revealed by our previous investigation of this region and nearby visual cortical areas (Tafazoli et al., 2017). Our work was motivated by the still limited and conflicting evidence about the involvement of these areas in advanced shape processing, which makes their assignment to a ventral-like stream much weaker, at the functional level, than one would expect on anatomical grounds (Gallardo et al., 1979; McDaniel et al., 1982; Wörtwein et al., 1993; Aggleton et al., 1997; Sánchez et al., 1997; Tees, 1999). For instance, while we observed an increase in the ability of rat lateral areas to support transformation-tolerant recognition, (Vermaercke et al., 2014) found that only TO (an area located laterally to LL) was superior to V1 and the other areas in supporting position tolerance, and only in relative terms – i.e., in terms of the stability, rather than of the magnitude, of the discrimination performance afforded by TO across two nearby positions. (Vermaercke et al., 2014) also found that orientation tuning increased along the areas’ progression – a trend that, as shown in our current study, is in disagreement with both the ventral stream literature and the behavior of ventral-stream models. In a later study, the same group reported a lack of object categorical representations in rat extrastriate lateral areas, which is unexpected for an object-processing stream (Vinken et al., 2016). At the same time, they found a decrease, along the lateral progression, in the accuracy of V1-like models to predict neuronal responses, as expected for a ventral-like pathway.

The literature probing the tuning properties of mouse extrastriate areas features a similar variety of findings. In mice, it is unclear whether an equivalent of rat area LL exists – earlier anatomical maps reported such area (Olavarria and Montero, 1989), although more recent studies would place the cortical field lateral to LI either in the anterior part of postrhinal cortex (PORa; also named posterior area 36) or in the posterior part of temporal cortex (TEp) (Wang and Burkhalter, 2007; Wang et al., 2012; Gămănuț et al., 2018). Regardless of the naming convention, this region has been attributed to the ventral stream on anatomical grounds (Wang et al., 2011, 2012), along with areas LM (lateromedial) and LI, which have been tested with drifting gratings in several electrophysiological and imaging studies. (Van den Bergh et al., 2010) measured the phase dependence of neuronal responses in V1 and nearby lateral cortex (named V2 by the authors, which, very likely, corresponds to area LM) and found that the latter had a lower proportion of simple cells (15% vs. 38% in V1). Conversely, the authors reported virtually no difference between the areas in terms of OSI. This result contrasts with the general increase of orientation tuning that was found in nearly all extrastriate areas (compared to V1) by (Marshel et al., 2011), although the increment of OSI was much larger in putative dorsal-stream areas than in LM and LI. A significant increase of OSI in LM, relative to V1, was also observed by (Tohmi et al., 2014), while other authors found virtually no differences between V1 and some of the putative dorsal- and ventral-stream areas (Andermann et al., 2011; Roth et al., 2012; Smith et al., 2017).

The variety and inconsistency of these findings motivated our further search for the signature of ventral processing in LL, where, to be fully complementary with our previous study, we did not probe neuronal representations using visual objects. With the goal of targeting those core tuning processes that are responsible for building up transformation tolerance and feature selectivity, we deployed drifting gratings and noise movies, and we compared the V1 and LL neuronal populations in terms of: 1) the phase-sensitivity of their responses; 2) the goodness of the RF structures recovered through STA; and 3) the sharpness and complexity (i.e., bimodality) of their tuning for orientation. This approach required deriving clear predictions about how these properties should evolve along a ventral-like pathway. This was achieved through a careful survey of the monkey literature and by simulating the basic computational architecture of the ventral stream using a conceptual model of ventral computations (HMAX) and a state-or-the-art deep convolutional network (VGG16). Because of the invariance-building operation, phase-sensitivity sharply decreased across HMAX layers (Fig. 4C). Concomitantly, because of the selectivity-building computation, orientation tuning became smaller (Fig. 7C), with the units acquiring preference for multiple orientations (Fig. 8C). The same drop of phase-sensitivity was observed in VGG16, where orientation tuning also decreased sharply (with a concomitant increase of bimodal tuning) in the last layers of the network (Fig. 11B-C). All these trends were matched by the empirical evidence gathered in our study, when comparing V1 to LL (Figs. 4A-B, 7A-B and 8A-B), thus suggesting that similar selectivity- and invariance-building computations are at work along the progression of lateral extrastriate areas that culminates with LL. The consistency among these trends and those observed across the primate ventral stream (Figs. 4D and 7D) adds further, critical support to this conclusion.

As such, our results strengthen our previous attribution of LL to a ventral-like stream (Tafazoli et al., 2017), while shedding new light on the possible origin of the discrepancies with and among earlier rodent studies (see previous paragraphs). For instance, while we presented very large (60°x110°), full-field gratings, (Vermaercke et al., 2014) used smaller (33°) circular gratings, shown at the RF center of each recorded neuron. Our presentation protocol ensured that, for most neurons in both areas, the stimuli covered the entire *classical* receptive field and, very likely, a large fraction of the *extra-classical* receptive field (Adesnik et al., 2012; Vaiceliunaite et al., 2013; Self et al., 2014; Pecka et al., 2014; Alwis et al., 2016), where surround-suppression can strongly affect shape tuning (Maunsell and Newsome, 1987; Orban, 2008). By contrast, the smaller and more localized gratings used in (Vermaercke et al., 2014) likely covered an increasingly smaller fraction of both the classical and extra-classical receptive fields while progressing from V1 to LL, given that RF size gradually increases along lateral extrastriate areas, being twice as large in LL as in V1 (Tafazoli et al., 2017). This, in turn, may have prevented the tuning for multiple oriented elements, located at different RF locations, to emerge in neurons with larger RFs, thus artificially boosting OSI in higher-order areas.

This interpretation is only partially supported by the monkey literature. In fact, although surround modulation has been shown to play a major role in boosting the selectivity of monkey visual neurons (Vinje and Gallant, 2000), most primate studies reviewed in Table 2 and Fig. 7D used circular gratings with a 6-12° diameter to probe orientation tuning, rather than full-field stimuli. Still, the size of those stimuli, relative to the typical RF size of ventral stream areas (i.e., median of 2° and 10°, respectively in V1 and IT), was slightly larger than the size of the circular gratings used by (Vermaercke et al., 2014), relative the median RF size of rat V1 and LL neurons (15° and 30° respectively), thus possibly engaging stronger extra-classical processing in monkey recordings. Alternatively, it is possible that in rats, even more than in monkeys, the signature of higher-order processing of shape information may clearly emerge only when surround-modulation mechanisms are fully engaged. Support to this conclusion comes from (Vinken et al., 2016), who, as previously mentioned, found a reduction in the ability of V1-like models (based on combinations of oriented Gabor filters) to predict neuronal responses to natural movies along the lateral progression. Not only this finding is in agreement with the poor estimates of the RF structures that we obtained in LL, as compared to V1, using STA, but strongly suggests that a reduction of orientation tuning has to be expected from V1 to downstream lateral areas (not tested in that study). That is, the conclusions of (Vinken et al., 2016), obtained with full-field stimuli (size 50°-74°), appear to be in better agreement with the drop of OSI reported in our study than with the increase of OSI previously found by the same group, using smaller, circular gratings (Vermaercke et al., 2014). To summarize, the discrepancies among the tuning properties of rat lateral neurons appear to be largely accounted for by differences in terms of visual stimuli, as argued here, as well as experimental design and data analysis, as discussed in our previous study (Tafazoli et al., 2017).

Similar arguments can be applied to interpret the results of mouse experiments, but only for the fraction of studies that used localized, circular gratings (Andermann et al., 2011; Smith et al., 2017) instead of full-field stimuli (Van den Bergh et al., 2010; Marshel et al., 2011; Tohmi et al., 2014). In the case of mouse studies, however, a more fundamental methodological difference exists with our and previous rat studies. In mice, many recordings were based on calcium imaging, performed using different indicators [e.g., OGB-1 (Marshel et al., 2011) vs. GCaMP3 (Andermann et al., 2011)], which diverge from a perfectly linear relationship between fluorescence changes and underlying spiking activity in an-indicator specific way (Akerboom et al., 2012). As pointed out by (Niell, 2011), this could potentially distort the orientation tuning curves and bias the OSI measurements, as compared to approaches where well-isolated single-units are recorded with micro-electrodes. In addition, while our recordings targeted cortical layer V (Fig. 1C), optical imaging studies typically target superficial layers. Although our previous study found the same proportional increase of ventral-specific processing from V1 to LL in both superficial and deep layers, the magnitude of this difference was smaller in the former (Tafazoli et al., 2017). This could make between-area differences harder to detect in superficial layers, especially when precise spike count estimates are not available, as in imaging studies. This argument does not apply to the experiments of (Van den Bergh et al., 2010), who performed single-electrode recordings. However, this study targeted only area LM, which has been reported to be functionally very similar to V1, never displaying any sign of ventral-like functional specialization (Andermann et al., 2011; Marshel et al., 2011; Tafazoli et al., 2017). Finally, it cannot be excluded the possibility that major differences exist, between mouse and rat visual cortex, in terms of high-order processing of shape information. In fact, despite some recent efforts (Aoki et al., 2017; Yu et al., 2018), mouse visual perception is still largely unexplored, and it remains unknown whether mice are able to perform those perceptual tasks, recently demonstrated in rats (Zoccolan et al., 2009; Tafazoli et al., 2012; Alemi-Neissi et al., 2013; Vinken et al., 2014; Rosselli et al., 2015; De Keyser et al., 2015; Djurdjevic et al., 2018), that should specifically engage the ventral stream.

To conclude, we believe that our results nicely complement and extend those of our previous study (Tafazoli et al., 2017). In fact, they show how some key neuronal tuning properties, which are revealing of the build-up of transformation tolerance and shape selectivity along an object-processing pathway, naturally evolve, from V1 to LL, in a direction that is consistent with the previously demonstrated ability of LL to support transformation-tolerant object recognition – an ability that, in rodents, is likely essential for ecologically relevant tasks such as spatial navigation and foraging (Cox, 2014; Zoccolan, 2015). In addition, being our experimental approach based on the presentation of widely-used parametric stimuli and the application of intuitive metrics, it provides a standardized and easily interpretable way to carry out a large-scale screening of ventral processing across rodent visual cortical areas. This could help new optical imaging studies finding the yet missing functional signature of ventral processing in mouse visual cortex – e.g., by analyzing existing large datasets of stimulus-response associations, as the one made available by the Allen Brain Observatory (Hawrylycz et al., 2016). In this regard, having provided clear predictions about the evolution of neuronal tuning along an object-processing pathway will likely serve as a reference to guide future explorations of ventral functions in both mice and rats.

## Acknowledgments

This work was supported by a European Research Council Consolidator Grant (DZ, project n. 616803-LEARN2SEE).

